# Targeted genotyping-by-sequencing of potato and data analysis with R/polyBreedR

**DOI:** 10.1101/2024.02.12.579978

**Authors:** Jeffrey B. Endelman, Moctar Kante, Hannele Lindqvist-Kreuze, Andrzej Kilian, Laura M. Shannon, Maria V. Caraza-Harter, Brieanne Vaillancourt, Kathrine Mailloux, John P. Hamilton, C. Robin Buell

## Abstract

Mid-density targeted genotyping-by-sequencing (GBS) combines trait-specific markers with thousands of genomic markers at an attractive price for linkage mapping and genomic selection. A 2.5K targeted GBS assay for potato was developed using the DArTag^TM^ technology and later expanded to 4K targets. Genomic markers were selected from the potato Infinium^TM^ SNP array to maximize genome coverage and polymorphism rates. The DArTag and SNP array platforms produced equivalent dendrograms in a test set of 298 tetraploid samples, and 83% of the common markers showed good quantitative agreement, with RMSE (root-mean-squared-error) less than 0.5. DArTag is suited for genomic selection candidates in the clonal evaluation trial, coupled with imputation to a higher density platform for the training population. Using the software polyBreedR, an R package for the manipulation and analysis of polyploid marker data, the RMSE for imputation by linkage analysis was 0.15 in a small half-diallel population (N=85), which was significantly lower than the RMSE of 0.42 with the Random Forest method. Regarding high-value traits, the DArTag markers for resistance to potato virus Y, golden cyst nematode, and potato wart appeared to track their targets successfully, as did multi-allelic markers for maturity and tuber shape. In summary, the potato DArTag assay is a transformative and publicly available technology for potato breeding and genetics.

**Core Ideas:**

1. A mid-density, targeted genotyping-by-sequencing (GBS) assay was developed for potato.
2. The GBS assay includes markers for resistance to potato virus Y, golden cyst nematode, and potato wart.
3. The GBS assay includes multi-allelic markers for potato maturity and tuber shape.
4. The polyBreedR software has functions for manipulating and imputing polyploid marker data in Variant Call Format.
5. Linkage Analysis was more accurate than the Random Forest method when imputing from 2K to 10K markers.

## 1 INTRODUCTION

Targeted genotyping-by-sequencing (GBS) has become an essential technology for molecular plant breeding. As with restriction site-associated DNA (RAD) sequencing (Baird et al., 2008; Elshire et al., 2011), targeted GBS is based on sequencing a reduced representation of the genome. A key difference is that targeted GBS uses a fixed set of primer pairs or oligonucleotide baits, with the number of targets designed based on the application and price point (Campbell et al., 2015; Gasc et al., 2016; Ali et al., 2016). DArTag is a targeted GBS method based on PCR with molecular inversion probes (Hardenbol et al. 2003) and scalable to thousands of targets (Hardigan et al., 2023; Zhao et al., 2023). As part of the CGIAR Excellence in Breeding platform, DArTag panels were developed for wheat, maize, rice, cowpea, pigeon pea, common bean, groundnut (peanut), sorghum, and potato (Excellence in Breeding, 2022). This article describes the design and validation of the first potato DArTag panel, which had 2.5K targets, as well as a second design project, which extended the assay to 4K targets.

Before DArTag, there was no comparable “mid-density” genotyping service for potato. The main genotyping platform for genetic mapping and genomic selection in potato has been an Infinium^TM^ SNP array, which was originally developed with 8303 markers and then expanded to 12K (Version 2) based on the same discovery panel of 6 varieties (Hamilton et al., 2011; Felcher et al., 2012). The 22K V3 array incorporated new SNPs from a larger discovery panel of 83 tetraploid varieties (Uitdewilligen et al., 2013; Vos et al., 2015), and the 31K V4 array added markers from yet another discovery pool (Sharma and Bryan, 2017). To maintain backwards compatibility with existing marker data, the genomic markers for DArTag were selected from the potato Infinium array.

In addition to genomic markers, the DArTag design includes “essential markers” that are prioritized during the final marker selection and primer design process. For potato, our initial priority was identifying markers with high diagnostic value (i.e., haplotype-specificity) for key resistance genes in a wide variety of genetic backgrounds. When the V1 assay was designed in 2020, KASP markers for the *Ry_adg_* (Herrera et al., 2018) and *Ry_sto_* (Nie et al., 2016) resistance genes against potato virus Y (PVY) were being widely utilized through a “low-density” genotyping service of the Excellence in Breeding platform. These two markers were therefore obvious candidates to include in the V1 DArTag panel. When the V2 DArTag panel was designed in 2023, a number of additional traits were targeted with essential markers.

R package polyBreedR (https://github.com/jendelman/polyBreedR) was initiated in 2020 to publicly distribute software used for genomics-assisted breeding of potato (Endelman et al. 2017; Endelman et al. 2018). Since elite potato germplasm is autotetraploid, all polyBreedR functions work for both diploid and tetraploid marker data, and most work for higher ploidy too. The focus of the original functions was SNP array data, which was more commonly used than GBS in potato and other polyploids due to the high read depth needed to differentiate heterozygotes with different allele dosage (Uitdewilligen et al., 2013). The required depth for < 5% error in a tetraploid ranges from 30 to 60, depending on the population structure and other assumptions (Gerard et al. 2018; Matias et al. 2019). Because the number of genome fragments is much smaller with mid-density, targeted GBS than RAD-seq, read depth is less of a limitation. In parallel to developing the DArTag assay for potato, new functions were added to polyBreedR to facilitate the manipulation and imputation of marker data in Variant Call Format (VCF).

## 2 MATERIALS AND METHODS

### 2.1 Genomic markers

SNPs were selected from the 22K V3 SNP array (Felcher et al., 2012; Vos et al., 2015) for the 2.5K V1 DArTag set, and additional SNPs were selected from the V4 31K SNP array for the 4K V2 DArTag set. Physical positions were based on the DMv6.1 reference genome (Pham et al., 2020). Genetic map positions (in cM) were interpolated from the map positions reported in Endelman and Jansky (2016). The interpolated “Marey” map of cM vs. bp was constrained to be monotone nondecreasing (Figure S1) using an I-spline basis with 12 degrees of freedom, generated with R/splines2 (Wang and Yan, 2021). Non-negative basis coefficients were computed by minimizing the mean-squared error with R/CVXR (Fu et al., 2020). The script is available as function *interpolate_cM* in R/MapRtools (Endelman, 2023a). Initially, SNPs were selected based on discretizing the genome into 1 cM bins, and within each bin, SNPs were prioritized based on minor allele frequency (MAF) in a collection of US and CIP germplasm. After saturating the genome, additional SNPs were selected sequentially based on the ad-hoc score *d* + 10 × *MAF*, where *d* is cM distance to the closest selected SNP.

Germplasm for evaluating V1 DArTag came from the International Potato Center (CIP) and University of Wisconsin breeding programs. Data for 703 tetraploid samples are provided as VCF File S1, with the year of submission for each sample recorded in File S2. The function *dart2vcf* in polyBreedR generates a VCFv4.3 compliant file from the two standard DArTag CSV files (“Allele_Dose_Report” and “Allele_match_counts_collapsed”). polyBreedR function *gbs* was used to replace the original DArTag genotype calls (FORMAT field GT) with those from function *flexdog* in R/updog, using the “norm” prior (Gerard et al., 2018). Three parameters of the beta-binomial model (SE = sequencing error, AB = allelic bias, OD = overdispersion) were stored for each variant. Both functions utilize R package vcfR (Knaus and Grunwald, 2017).

One submission of tetraploid (N=323) and diploid (N=52) samples was used to evaluate the 4K V2 DArTag assay. VCF File S3 contains the target SNP data, and CSV File S4 is the MADC (“missing allele discovery counts”) file from DArT, which contains read counts for 81bp haplotypes (similar to a TagsByTaxa file in TASSEL; Glaubitz et al. 2014). Genotype calls were made separately for each ploidy group using the *gbs* function, and the resulting VCF files were then combined with *bcftools merge* (Danecek et al. 2021); the model parameters in INFO are from the tetraploid set. To verify ploidy, genotype calls were made for the diploid samples based on the tetraploid model parameters, using the “model.fit=FALSE” option of function *gbs*, which in turn uses *flexdog* with “update_bias”, “update_seq”, and “update_od” set to FALSE.

A comparison of DArTag vs. SNP array genotypes was conducted using 298 clones for V1 DArTag and 78 clones for V2 DArTag. XY intensity values and genotype calls are provided in File S5 for 15,187 markers from the V4 SNP array, based on a normal mixture model estimated with R/fitPoly (Voorrips et al., 2011; Zych et al., 2019). The parameter file for the normal mixture model is distributed with the polyBreedR package as “potato_V4array_model.csv” and was used to convert Genome Studio Final Reports to VCF with function *array2vcf*. This function also requires a VCF map definition file to convert from B allele dosage to ALT dosage, which is distributed as “potato_V4array.vcf” with polyBreedR. The common markers between DArTag and the SNP array, including matching REF/ALT, were identified using *bcftools isec*.

### 2.2 Imputation

Two methods were compared for the accuracy of imputing SNP array markers from DArTag: Random Forest (RF) and Linkage Analysis (LA). Method RF was implemented as polyBreedR function *impute_L2H*, using the R/randomForest package (Liaw and Wiener, 2002). The number of trees was set at 100 by monitoring the out-of-bag error, using the default number of variables randomly sampled at each split, which is √*m* for *m* classification variables. Each marker was imputed separately, using the *m* closest markers as candidate prediction variables.

Results were generated for *m*=10, 25, 50, and 100, and the lowest error (Table 1) was observed at 25, but the optimal marker number will vary by dataset. Method LA was implemented as polyBreedR function *impute_LA*, using the software PolyOrigin (Zheng et al., 2021) and default parameters. Imputation error was measured using leave-one-family-out cross-validation in a five-parent half-diallel population (pedigree in File S6). The parent codes in Table 1 are P1=W6609-3, P2=W12078-76, P3=W13NYP102-7, P4=W14NYQ4-1, P5=W14NYQ9-2. The high density (10K) phased parental genotypes are in File S7.

**Table 1.**
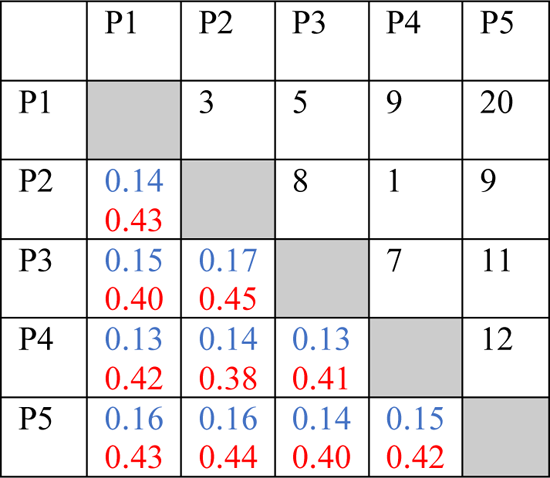
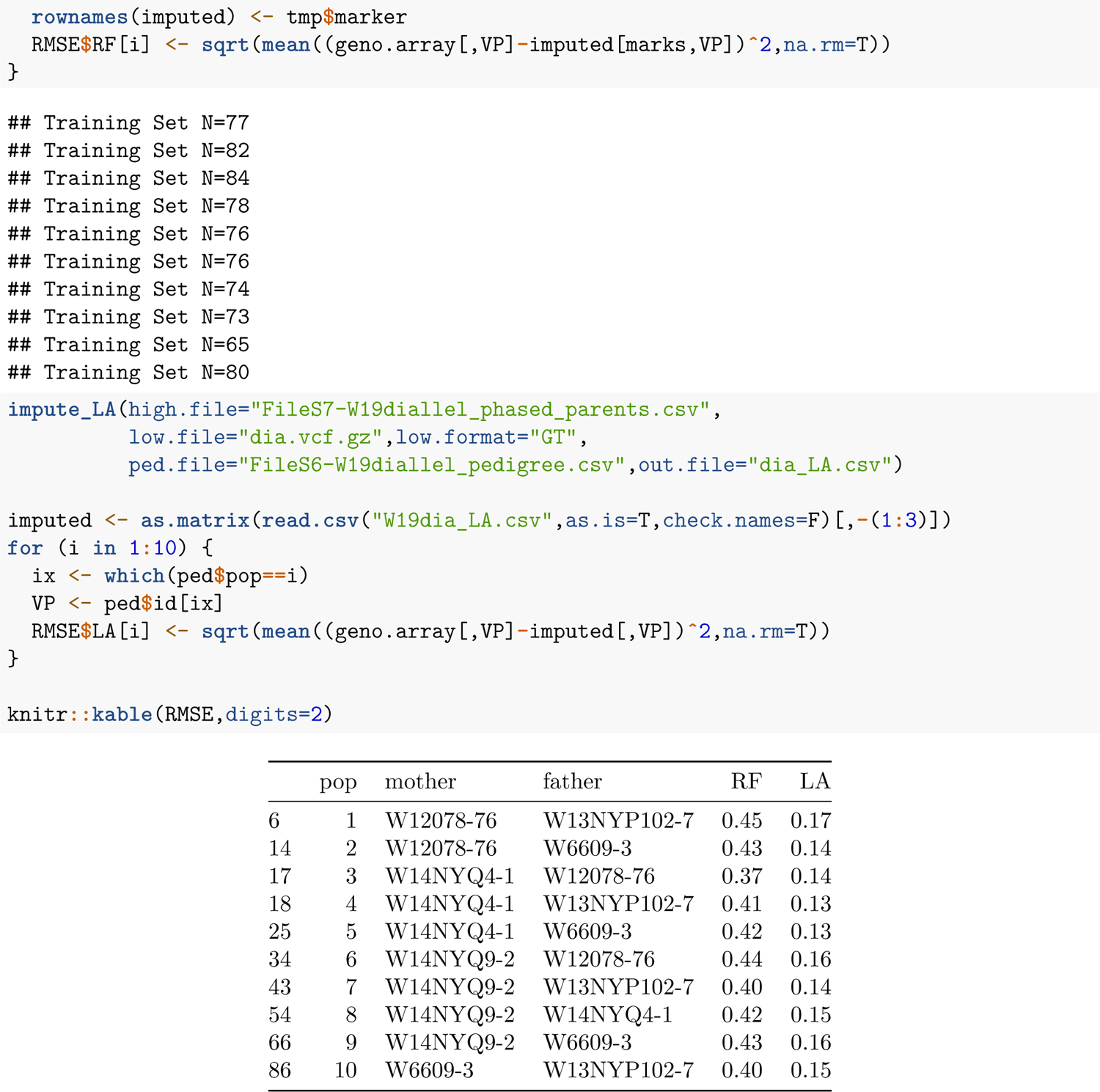
Half-diallel population with five parents. Above diagonal: F1 population sizes; Below diagonal: imputation root-mean-squared-error with linkage analysis (blue, top) vs. random forest (red, bottom).

### 2.3 Trait markers

A set of six interconnected F1 populations was used to assess the accuracy of genotype calls for the V2 DArTag trait markers Ryadg_chr11_2499502 and H1_chr05_52349069. Parental phasing and haplotype reconstruction utilized PolyOrigin (Zheng et al., 2021), and binary trait locus (BTL) analysis utilized R/diaQTL (Amadeu et al., 2021). The pedigree, genomic marker, and dominant trait marker files needed for diaQTL are Files S8, S9, S10, respectively. Validation of trait marker Sli_chr12_2372490 was based on the Sli_898 KASP marker (Clot et al., 2020; Kaiser et al., 2021).

Trait markers CDF1.2_chr05_4488015 and CDF1.4_chr05_4488021 target two different 7 bp insertions of *CDF1* (Kloosterman et al., 2013; Gutaker et al., 2019) and contain equivalent information in the DArT MADC file. The 81-bp haplotypes were aligned using MUSCLE v3.8 (Edgar, 2004). DArTag read counts for *CDF1* alleles 1, 2, and 4 were tabulated with polyBreedR function *madc* and validated against genotypes determined via whole-genome sequencing with NovaSeq 2×150 reads (Song and Endelman, 2023).

Genome assemblies of *S. tuberosum* dihaploids were used to validate markers for *OFP20*, a major gene affecting tuber shape (Wu et al. 2018). High molecular weight DNA was extracted from tissue culture plantlets using a CTAB isolation method and Qiagen Genomic tips (Hilden, Germany), followed by an Amicon filter (MilliporeSigma, Burlington, MA) buffer exchange (Vaillancourt et al., 2019) or Takara NucleoBond HMW DNA kit (Takara, Kusatsu, Shiga, Japan). Genome assembly used hifiasm v0.16.1-r375 (Cheng et al. 2021, 2022) with PacBio HiFi Sequel II (Menlo Park, CA) reads from the University of Minnesota Genomics Center. Contigs less than 50kb were discarded using seqkit v2.3.0 (Shen et al. 2016), followed by Ragtag v2.1.0 (Alonge et al. 2019) to scaffold with DM 1-3 516 R44 v6.1 (Pham et al., 2020).

A multiple sequence alignment of 19 *OFP20* haplotypes (File S11) was generated using MUSCLE v3.8. Alleles 1–7 and M6_ScOFP20 were reported by van Eck et al. (2022), and the remaining haplotypes come from the dihaploids. The frequency of *OFP20.1* was approximated by ALT frequency at marker OFP20_M6_CDS_994 (994 bp in M6 CDS). For allele *OFP20.8,* which was discovered in the dihaploids (i.e., not in the FASTA file from van Eck et al. (2022)), allele frequency was approximated by REF frequency at marker OFP20_M6_CDS_24; this only works in populations without the M6_ScOFP20 allele. Marker OFP20_M6_CDS_171 was used to report allele depth for allele 2 (ALT) vs. alleles 3 and 7 combined (REF); alleles 1 and 8 were not detected by this marker. Marker OFP20_M6_CDS_75 was supposed to capture an indel at 82 bp that differentiates alleles 3 and 7, but neither haplotype was present in the MADC File S4.

## 3 RESULTS

### 3.1 Genomic markers

Version 1 (V1) of the potato DArTag GBS assay contained 2501 genomic SNPs, which were selected from the 22K V3 potato SNP array to maximize genome coverage and polymorphism rates (i.e., high minor allele frequency). The number of genomic markers per chromosome ranged from 176 on chr12 to 272 on chr01. The mean distance between adjacent markers was 0.35 cM, with the largest gap of 4.77 cM located on chr11 (Figure S2).

Analysis of 703 tetraploid samples, from three submissions across three years (2020-2022), revealed variability in the amount of sequencing data per sample. In 2020, the total depth (DP sum over markers) was consistent across samples, with mean 0.53M/sample and standard deviation 0.07M (Figure 1). The distribution in 2021 was bimodal, with the two modes corresponding to different plates. The lower mode was 0.63M, while the higher mode was 0.96M. The average total depth in 2022 was similar to 2020, at 0.53M/sample, but the standard deviation was higher, at 0.17M.

**Figure 1.**
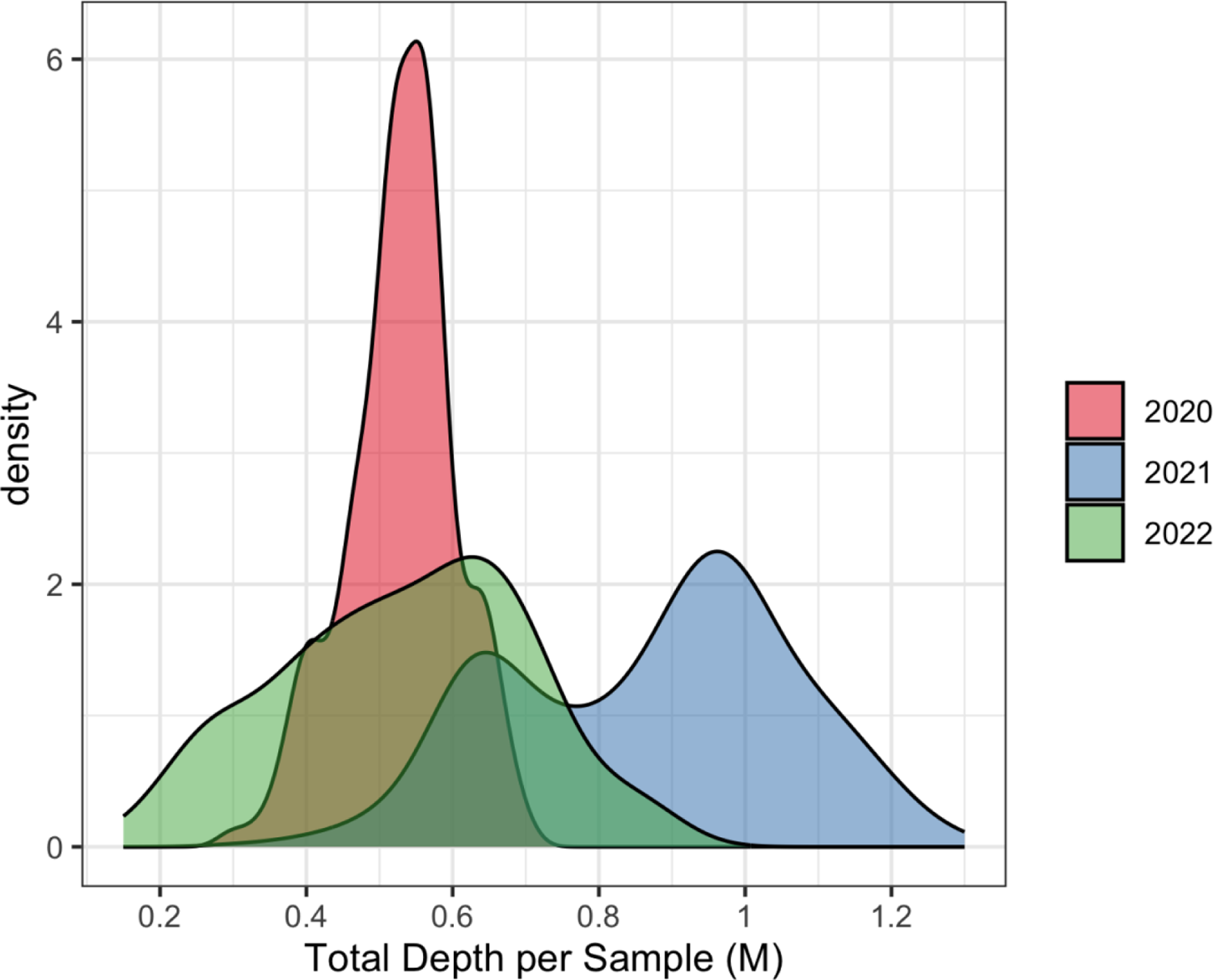
Total depth per sample, in million (M) read counts, for three submissions of potato V1 DArTag.

When sample DP was summarized by marker, the data were more consistent across years (Figure 2). The 10^th^ percentile for mean sample DP was 32, 53, and 24 in years 2020, 2021, and 2022, respectively (Fig. 2A). Despite the observed differences in total DP per sample (Figure 1), there was a consistent relationship between the mean (μ) and standard deviation (σ) for sample DP (Fig. 2B). The relationship between these quantities in a Poisson distribution is μ = σ^0.5^, which is a straight line with slope 0.5 on a log-log plot (dashed line in Fig. 2B). The observed data were overdispersed (i.e., more variable) compared to the Poisson, with slope 0.79 (SE 0.00), meaning that μ ≈ σ^0.8^.

**Figure 2.**
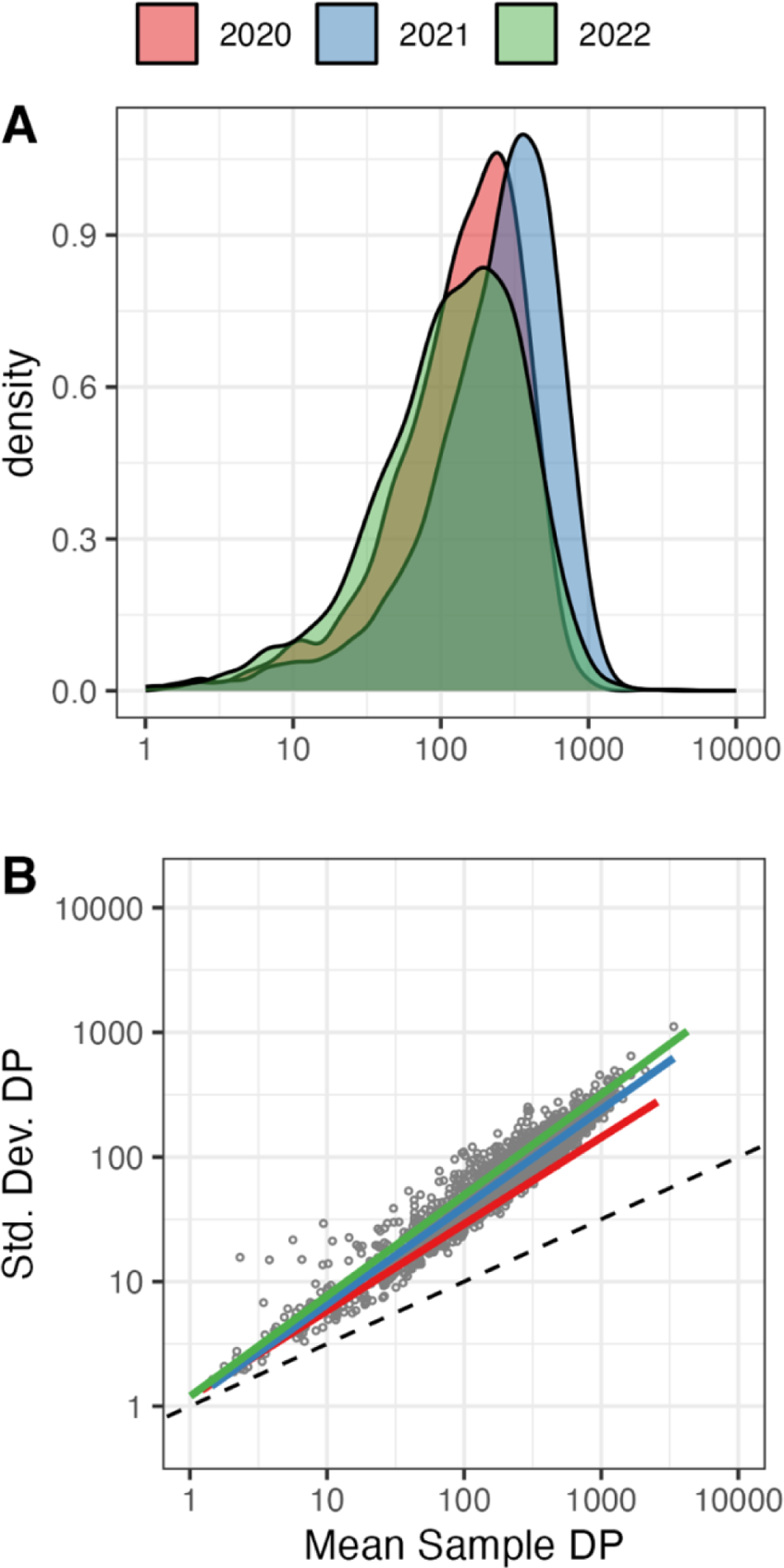
(A) Distribution of the mean sample depth (DP) for V1 DArTag markers. (B) Log-log plot of the relationship between the standard deviation and mean for sample DP. Individual marker points are shown only for 2021 to maintain legibility. Combining the data across years, the overall regression line (not shown) has slope 0.79 (SE 0.00) and R^2^ = 0.99.

Tetraploid genotype calls were made with R package *updog* (Gerard et al. 2018), which provides estimates of allelic bias (AB) for each marker—a parameter that measures the relative probability of observing the REF vs. ALT allele. When AB=1, or equivalently log_2_(AB) = 0, there is no bias. When AB=2, or equivalently log_2_(AB)=1, the REF allele is twice as likely to be observed in a balanced heterozygote. 10% of the markers exhibited bias |(AB) | > 1 (Figure S3), but many of these still appeared to have reliable clustering (Figure 3).

**Figure 3.**
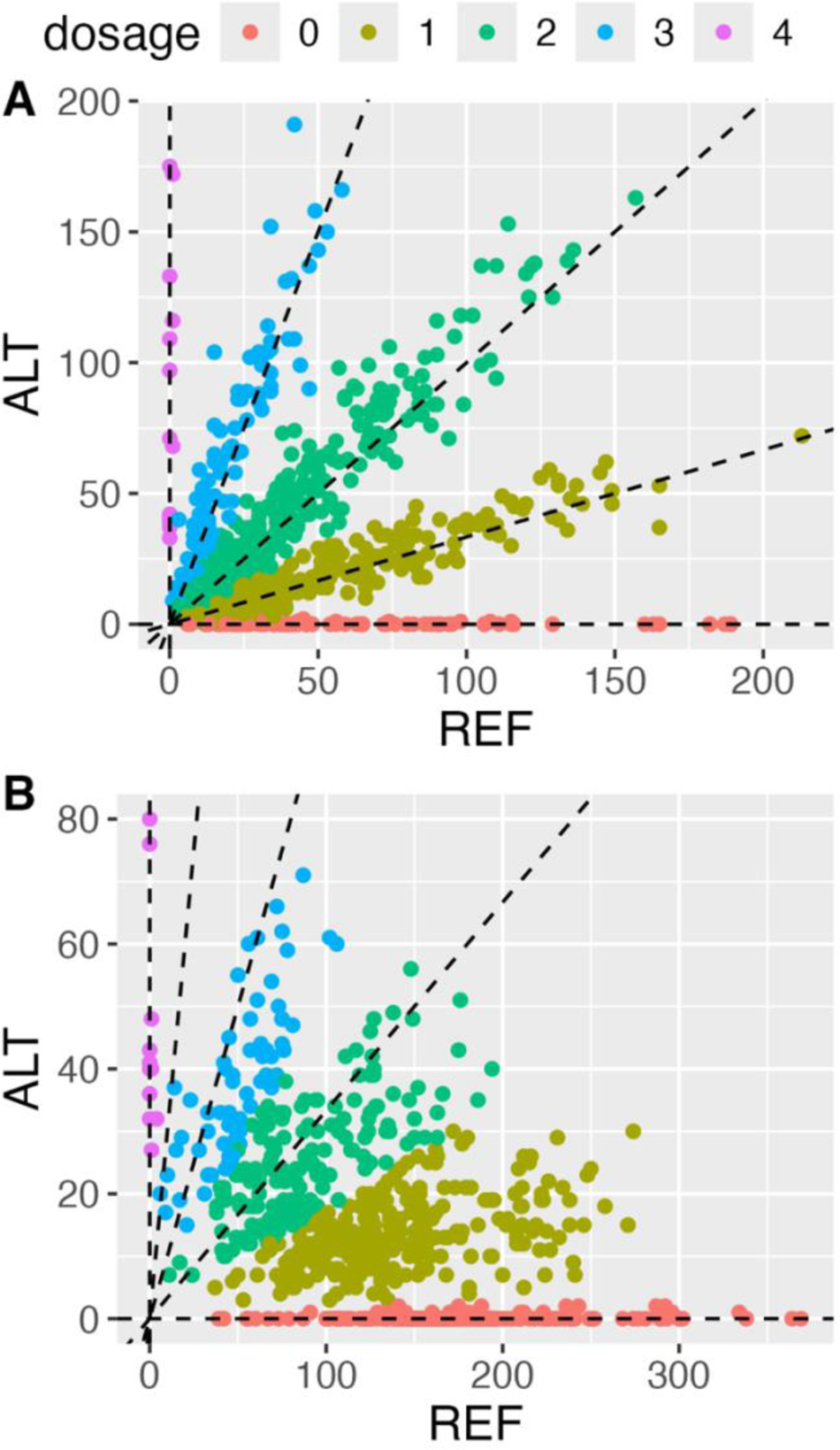
Examples of DArTag markers without (A) vs. with (B) allelic bias. Dashed lines correspond to possible tetraploid allele ratios when there is no allelic bias (1:0, 3:1, 1:1, 1:3, 0:1). (A) solcap_snp_c2_36615 with bias = −0.2. (B) PotVar0072076 with bias = 1.8.

V1 DArTag and SNP array genotypes were compared for 1865 common markers across 298 tetraploid clones. Both platforms identified two groups of genetically identical clones, one pair and one threesome, originating from the same F1 populations (Figure S4). This is not uncommon in potato breeding due to how single plant selection is conducted in the first field year. After removing duplicates, the two marker profiles (GBS & array) for every clone were paired under hierarchical clustering (Figure S5), indicating close agreement.

For a quantitative comparison, several measures of error were computed for each marker (File S12). Classification error (CE), which is the proportion of samples with different genotype calls, was calculated for both tetraploid (4x) and pseudo-diploid (2x) genotypes (where differences in heterozygote allele dosage are ignored). There was a sharp bend in the cumulative distribution for 2x CE at approximately 0.1 error (Figure 4), with 1647 markers below this threshold (88% of those tested). As expected, fewer markers (1302) satisfied 4x CE < 0.1 because of the difficulty discriminating between heterozygous genotypes. For 4x genotypes, the root-mean-squared-error (RMSE) of allele dosage is potentially more meaningful than CE, and 1547 markers had RMSE < 0.5 (Figure 4), a somewhat arbitrary threshold selected because it represents the midpoint between integer dosages.

**Figure 4.**
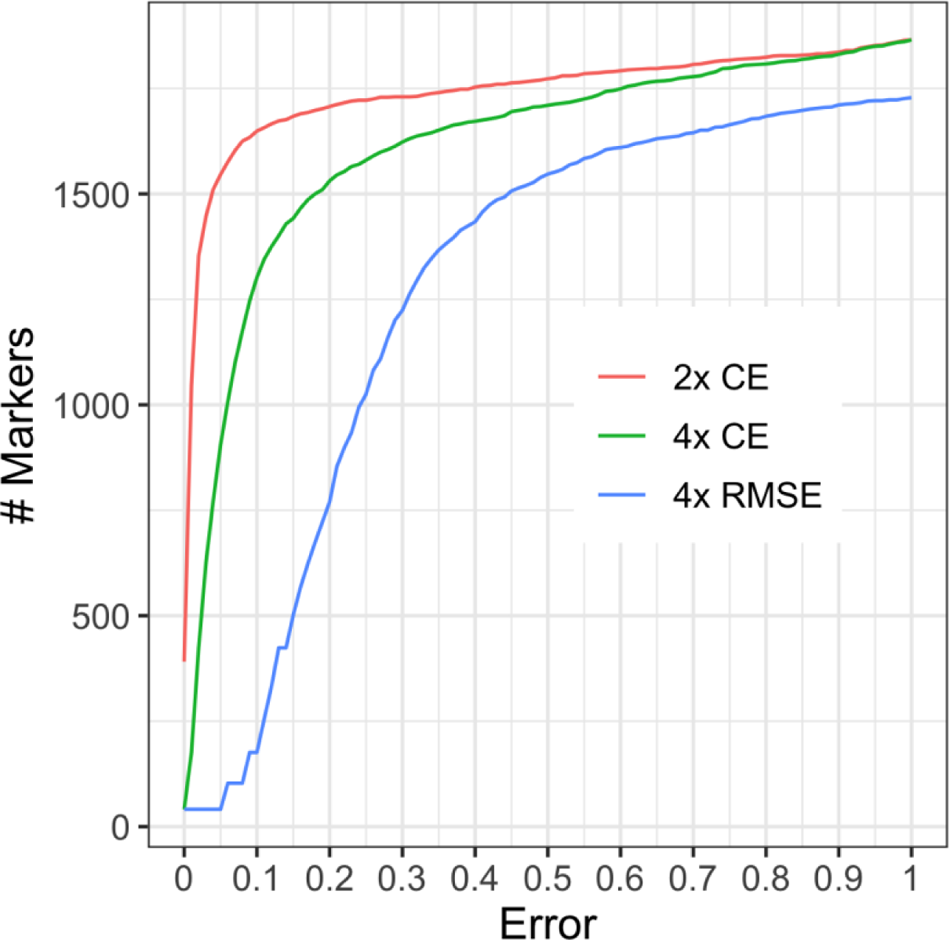
Empirical cumulative distribution for the error between the V1 DArTag and SNP array on 1865 common markers. CE = classification error. RMSE = root-mean-squared-error. 2x = pseudo-diploid genotypes. 4x = tetraploid genotypes.

Version 2 (V2) of the potato DArTag GBS assay was designed in 2023 and contains 3893 genomic SNPs, of which 2144 were included in V1. The additional SNPs were selected from the 31K V4 potato SNP array using the same criteria as before. GBS and SNP array genotypes were compared for 2608 common markers across 78 clones (40 tetraploid, 38 diploid). Given the small number of tetraploids, only the 2x CE criterion was computed, and 2341 markers had 2x CE < 0.1 (Figure S6; File S13).

### 3.2 Imputation and ploidy analysis

A key role for the DArTag genomic markers is to facilitate imputation to higher density platforms for genomic selection. Among the 298 clones genotyped with both the SNP array and V1 DArTag is a five-parent half-diallel population of 85 clones, with F1 family sizes between 1 and 20 (Table 1). The accuracy of two imputation methods was compared by leave-one-family-out cross-validation: Random Forest (RF) vs. Linkage Analysis (LA). Linkage analysis uses a genetic model of recombination and phased parental genotypes to reconstruct progeny in terms of parental haplotypes. The RMSE for imputing 10K SNP array genotypes from DArTag was always lower with LA compared to RF (Table 1), with overall means of 0.15 and 0.42, respectively.

The *check_ploidy* function of polyBreedR was originally developed to use SNP array markers to differentiate diploid from tetraploid samples based on the following principle: when a bi-allelic genotype model developed for tetraploids is applied to a diploid sample, ideally there would be no simplex (AAAB) or triplex (ABBB) heterozygotes, only duplex (AABB) calls because both AB and AABB genotypes have 1:1 allele ratios. The proportion of calls that are simplex or triplex can therefore be used to discriminate ploidies. This method has been used extensively during recent haploid induction crosses of tetraploid potato, where the desired result is a diploid haploid, aka dihaploid, containing only the maternal chromosomes, but sometimes the progeny are tetraploid or aneuploid (Amundson et al. 2020; Busse et al. 2021).

The *check_ploidy* function was extended to allow VCF file input and the R/updog model for genotype calling of GBS markers. When applied to the V2 DArTag dataset (File S3), the 323 tetraploids were clearly separated from the 52 diploids (Figure 5).

**Figure 5.**
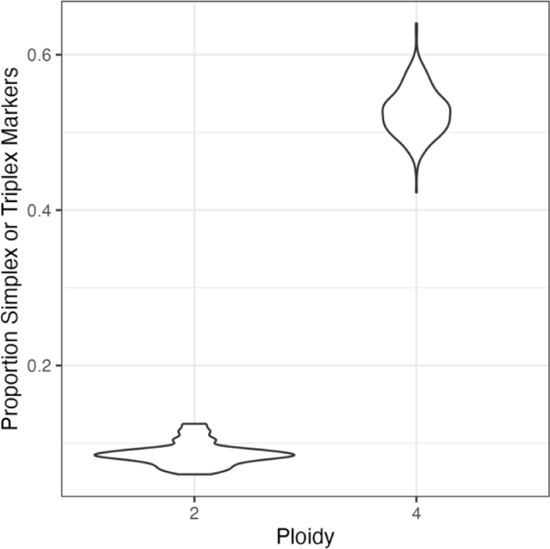
Violin plot illustrating ploidy discrimination with the V2 DArTag assay, based on the proportion of simplex or triplex markers with a tetraploid genotype calling model. The diploid potato samples had much lower values of this parameter.

### 3.3 Trait markers

The V1 DArTag assay had only two trait markers, targeting two different resistance genes (*Ry_adg_*, *Ry_sto_*) for potato virus Y (PVY), which is the most economically important viral pathogen of potato. Both variants had previously been targeted with KASP markers, and for 93 samples genotyped with both KASP and V1 DArTag, there were 2 discrepancies for presence/absence of *Ry_adg_* (Table S1). Both PVY markers were carried forward to the V2 DArTag assay, and four clones tested positive for *Ry_sto_*: three were expected based on previous testing, and the fourth was plausible based on its pedigree (Table S2). Many samples in the V2 submission tested positive for *Ry_adg_*, which was expected given its high frequency in US chip processing germplasm, but the allele dosages seemed too high—eight samples were even homozygous tetraploids. To investigate further, a partial diallel population (N=123) within the dataset was analyzed (Figure S7). Treating the *Ry_adg_* marker as a dominant trait, joint linkage analysis identified which parental haplotypes carry the R gene (Figure S8), and corrected dosages were determined by reconstructing the progeny in terms of parental haplotypes (Figure 6). Five triplex and two quadriplex calls for *Ry_adg_* were corrected down to duplex, and the average upward bias was 0.24 dosage.

**Figure 6.**
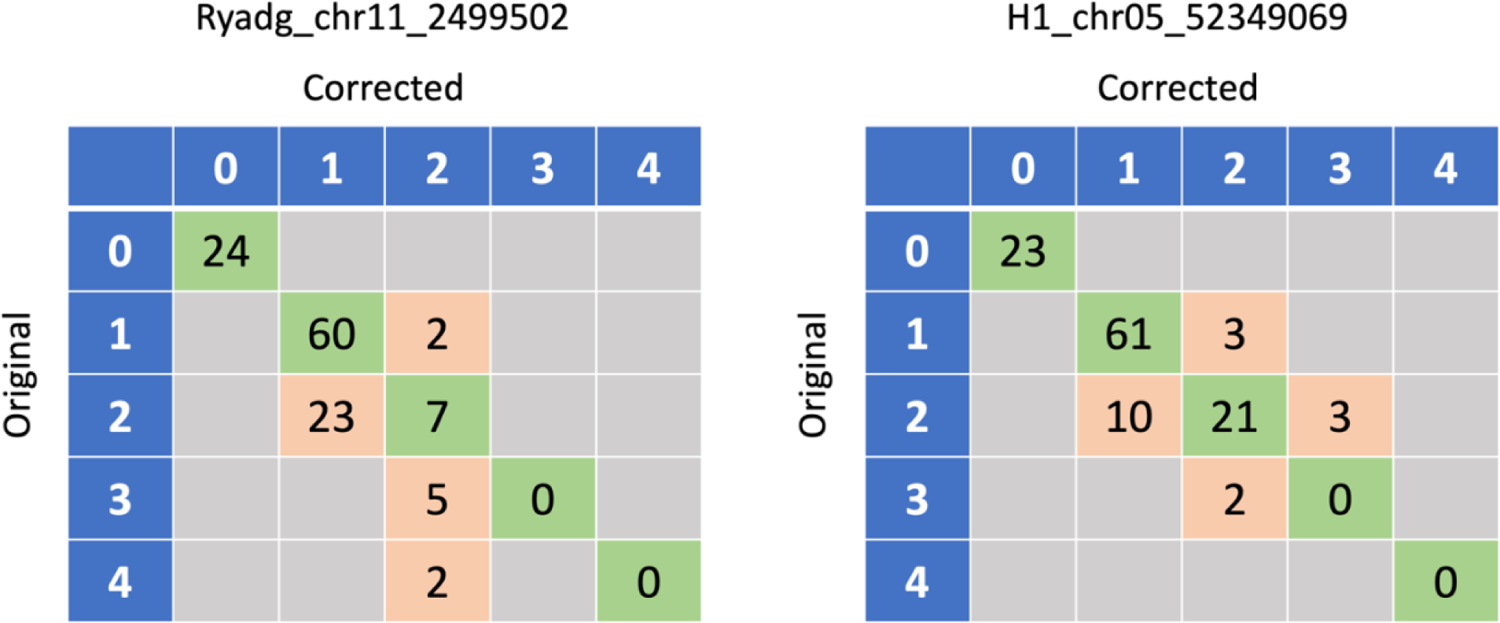
Original vs. corrected genotypes for the trait markers Ryadg_chr11_2499502 and H1_chr05_52349069. The original genotypes were based on R/updog with a “norm” prior and then corrected based on linkage analysis.

Besides the two PVY markers, the V2 DArTag assay has five additional trait markers with reliable results (Table 2). Like *Ry_adg_*, the golden cyst nematode resistance gene *H1* was common in US chip processing germplasm, but diallel analysis indicated the *H1* marker calls were more accurate, with an average bias of only 0.05 dosage (Figure 6).

**Table 2.** Validated trait markers in the V2 DArTag assay.

| Marker | Target Gene (Trait) | Functional Allele | Reference |
| --- | --- | --- | --- |
| Rysto_chr12_2352742 | *Rysto* (PVY) | REF<sup>a</sup> | Nie et al. (2016) |
| Ryadg_chr11_2499502 | *Ryadg* (PVY) | ALT | Herrera et al. (2018) |
| H1_chr05_52349069 | *H1* (golden cyst nematode) | ALT | Meade et al. (2020) |
| Sen3_chr11_2563398 | *Sen3* (wart) | ALT | Prodhomme et al. (2019) |
| Sli_chr12_2372490 | *Sli* (self-compatibility) | ALT | Clot et al. (2020) |
| CDF1.4_chr05_4488021 | *CDF1* (maturity) | N/A | Gutaker et al. (2019) |
| OFP20_M6_CDS_24<br>OFP20_M6_CDS_171<br>OFP20_M6_CDS_994 | *OFP20* (tuber shape) | N/A | van Eck et al. (2022) |

<sup>a</sup> The REF/ALT designation for Rysto\_chr12\_2352742 is reversed compared to the original design file based on the DMv6.1 reference genome. As a result, the functional allele is REF.

Little is known about resistance to potato wart disease (*S. endobioticum*) in US germplasm, but given the prevalence of the disease in other parts of the world (Obidiegwu et al., 2014), it has become a higher priority for molecular breeding. One trait marker targets the *Sen3* resistance gene, which was detected in four individuals with a common parent, AW07791-2rus. Based on pedigree information, the resistance appears to have been inherited from the maternal parent, PALB0303-1 (Elison et al., 2021).

Another trait marker targets *Sli*, a non-S locus F-box protein that disrupts the gametophytic incompatibility system and allows for the development of diploid, inbred lines (Ma et al., 2022; Eggers et al., 2022). The GBS marker showed perfect agreement with prior knowledge for 28 diploid samples based on KASP marker screening (Table S3).

A trait marker for the maturity gene *CDF1* targets the location of the 7 bp indel variants that differentiate alleles 2 and 4 from wild-type alleles, collectively designated group 1. Because of the multi-allelic nature of this variant, correct interpretation requires use of the DArT “missing allele discovery count” (MADC) file, which contains read counts for 81 bp haplotypes surrounding each target variant. Five CDF1 haplotypes were detected in the population (Figure 7A): three were full-length variants of CDF1.1 (Ref, Other1, Other2), one was CDF1.4 (Alt), and one was CDF1.2 (Other3). The validity of the assay was confirmed by comparing the read counts with samples of known CDF1 genotype (Figure 7B), with the complication that CDF1.3, which has an 865 bp transposon insertion at the same position, is not detected. As a result, samples with zero (or near zero, due to sequencing error) counts are interpreted as homozygous for allele 3. And since clones selected under long-day conditions are typically not homozygous wild-type, when CDF1.1 alleles are detected but not alleles 2 or 4, the predicted genotype is 1/3.

**Figure 7.**
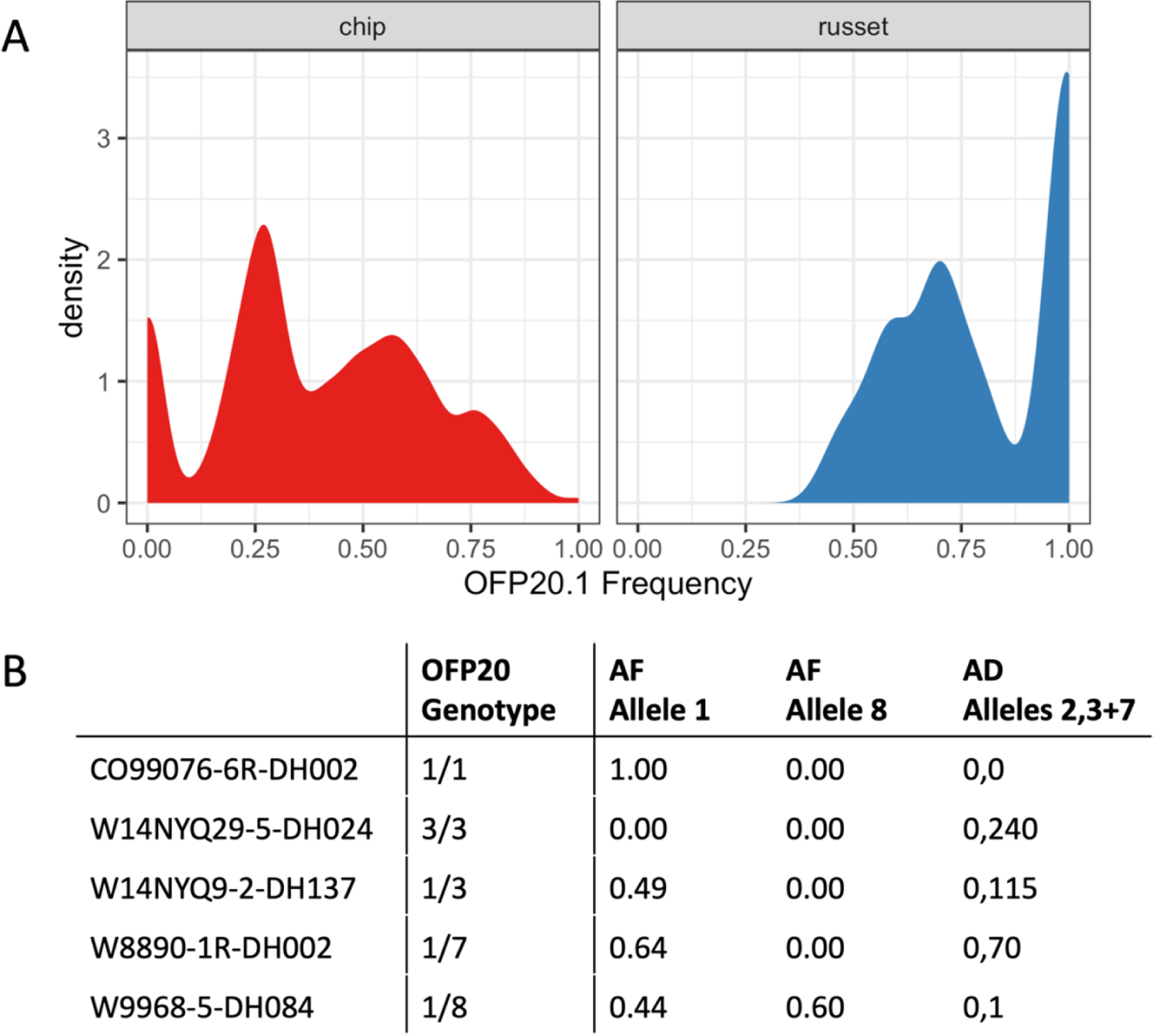
(A) Multiple sequence alignment of the DArTag haplotypes discovered for trait marker CDF1.4_chr05_448021. Haplotypes Ref, Other1, Other2 are CDF1.1 alleles, while Alt is CDF1.4 and Other3 is CDF1.2. (B) Haplotype read counts for samples with known CDF1 genotype.

Several markers were included in the V2 panel to target *OFP20*, an ovate family protein with a major effect on tuber shape (Wu et al. 2018). This is a complex locus with dozens of predicted alleles (van Eck et al. 2022), so the following approach to interpreting the DArTag markers may not work in all germplasm groups. Marker OFP20_M6_CDS_994 was used to estimate the frequency of *OFP20.1*, which is the most common allele in cultivated germplasm and promotes elongated shape (van Eck et al. 2022). *OFP20.1* was present at a higher frequency in the russet (N=21) vs. chip (N=300) samples from the UW breeding program (Fig. 8A), which is consistent with the long vs. round tuber phenotypes required for those market types. Marker OFP20_M6_CDS_24 was used to estimate the frequency of *OFP20.8*, which was present in 13% of the chip samples. Together with OFP20_M6_CDS_171, which provided information about presence/absence of *OFP20* alleles 2, 3, and 7, the DArTag markers were able to correctly predict five different *OFP20* genotypes (Fig. 8B).

**Figure 8.**
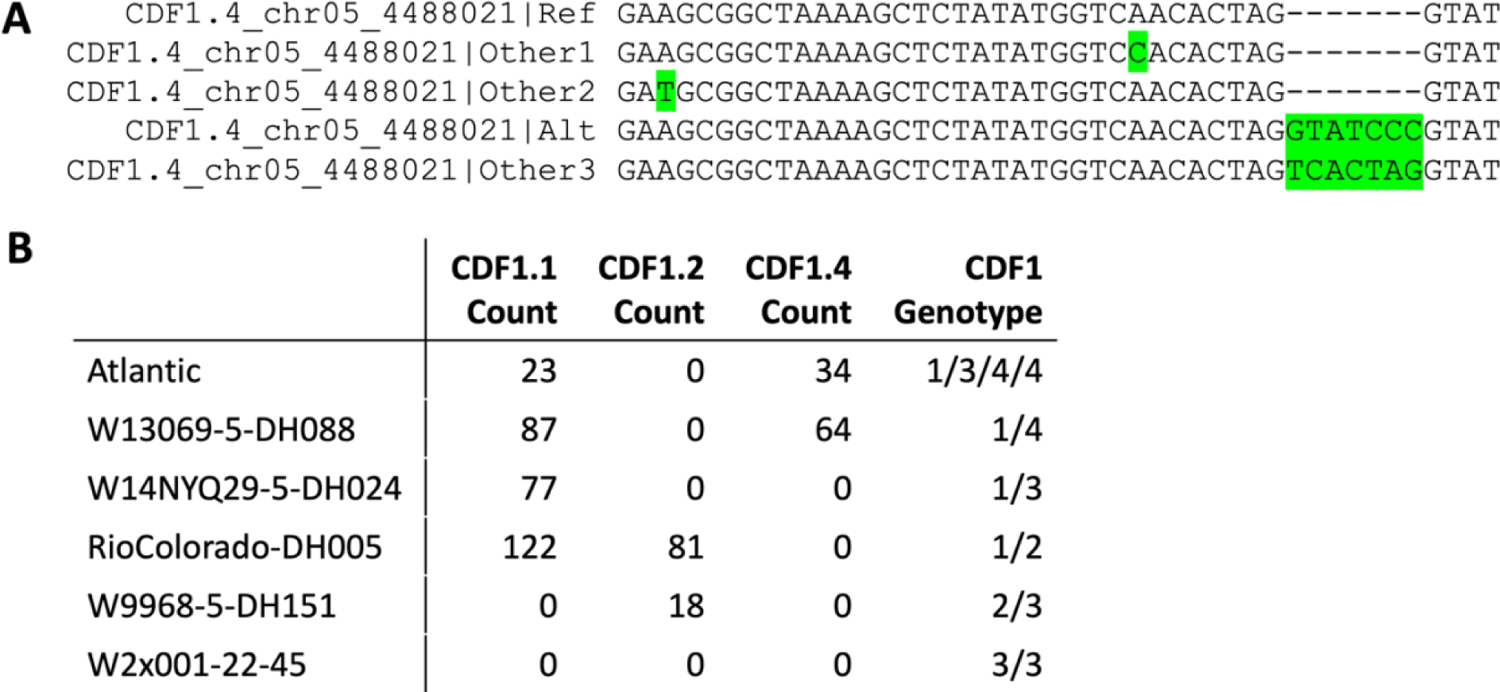
(A) Distribution of sample allele frequencies for OFP20.1 in round chip (N=300) vs. long russet (N=21) germplasm. (B) Comparison of known OFP20 genotypes with V2 DArT markers. Allele frequency (AF) of OFP20.1 was approximated by ALT frequency at marker OFP20_M6_CDS_994. AF of OFP20.8 was approximated by REF frequency at marker OFP20_M6_CDS_24. Allele depth (AD) at OFP20_M6_CDS_171 was used to distinguish allele 2 (ALT) from alleles 3 and 7 (REF).

## 4 DISCUSSION

The potato DArTag assay has several applications in potato breeding. For its price point, an ideal stage of deployment is the first clonal evaluation trial (CET), which typically occurs in the second field year of potato breeding and may have several thousand clones. The DArTag genomic markers provide a genetic fingerprint that can be used to correct pedigree errors (Muñoz et al., 2014; Endelman et al., 2017) and provide a reference genotype for quality control. The clonal trial entries are also candidates for genomic selection, both as potential clonal varieties and as parents to begin the next breeding cycle (Slater et al., 2016; Wu et al., 2023). Limited phenotyping for some traits occurs in the CET, and a genomic relationship matrix computed from DArTag markers could enable a multi-location trial to better estimate genetic values for the target population of environments, i.e., “sparse testing” (Endelman et al. 2014; Jarquin et al. 2020).

Based on previous studies, higher selection accuracy is expected if DArTag markers are first imputed to higher density (Cleveland and Hickey, 2013; Gorjanc et al., 2017). The exploitation of pedigree or family structure during marker imputation in diploids is well documented, with a range of methods and software available depending on the structure of the dataset (Meuwissen and Goddard, 2010; Swarts et al., 2014; Hickey et al., 2015; Whalen et al., 2018; Whalen et al., 2020). The present study has confirmed our hypothesis that linkage analysis is also beneficial for imputation in autopolyploids. DArTag panels are available for several autopolyploid crops besides potato, including alfalfa, blueberry, and sweetpotato (Breeding Insight, 2023), so the software developed for this study should benefit other breeding communities. Based on the current functionality of the PolyOrigin software (Zheng et al., 2021), only bi-allelic SNPs were used for imputation, but the DArTag MADC file offers the possibility of using multi-allelic haplotypes as markers, which are generally more informative for linkage analysis (Luo et al., 2001).

Besides more genomic markers, a major advantage of V2 compared to V1 DArTag is the additional trait markers (Table 2). It is very valuable to select for resistance to three important pests of potato—PVY, wart, and golden cyst nematode—with the same assay used for genomic selection. Notably absent from this list is potato late blight, caused by the pathogen *P. infestans*. Trait marker blb1_chr08_51070621 was designed to target the *RB/Rpi-blb1* gene (Song et al., 2003; van der Vossen et al., 2003) based on a SNP in the 3’UTR that worked well as a KASP marker (Sorensen et al., 2023). However, no haplotypes were detected in the V2 DArTag experiment for three positive samples from the KASP study. The V2 assay also targeted two genes affecting tuber skin color: *f3’5’h* (Jung et al., 2005) and *an2* (Jung et al., 2009). Both loci have complex allelic series (Hoopes et al. 2022), and more information is needed about their functional effects to guide selection. For tuber shape, a trait marker for the most common long allele (*OFP20.1*) can have an immediate impact on parent selection in the russet market type, where round alleles are undesirable due to their partial dominance. Table 3 summarizes the new or updated functions in polyBreedR associated with this research. Examples of their usage on a sample dataset can be found in Vignette 3 of the package and the Supplemental Methods file, which provides code to generate the main figures and tables.

**Table 3.** New or updated functions in R/polyBreedR developed for this research.

| Function | Description |
| --- | --- |
| *dart2vcf* | converts DArTag data files to VCF |
| *array2vcf* | converts Genome Studio Final Report to VCF file |
| *gbs* | adds genotype calls to a VCF file based on the R/updog model |
| *impute_L2H* | imputes low to high density markers by Random Forest |
| *impute_LA* | imputes low to high density markers by Linkage Analysis |
| *check_ploidy* | discriminates diploid from tetraploid samples |

## SUPPLEMENTAL FILES

During peer review, the supplemental files are available from the Dryad Digital Repository at https://datadryad.org/stash/share/tdvUt18gBCz6bJ568DaE7mLrWtq55kZzJa_C1uLYSfQ. The permanent link for the supplemental files after publication is https://doi.org/10.5061/dryad.8pk0p2nw4.

File S1. Potato DArTag V1 data for 703 samples (VCF).

File S2. Metadata with year submission for the samples in File S1 (CSV).

File S3. Potato DArTag V2 data for 375 samples (VCF).

File S4. DArT Missing Allele Discovery Counts for the samples in File S3 (CSV).

File S5. Potato V4 SNP array data for 298 samples (VCF).

File S6. Pedigree for diallel population in the V1 DArTag dataset (CSV).

File S7. Phased parental genotypes for the diallel population in File S6 (CSV).

File S8. Pedigree for diallel population in the V2 DArTag dataset (CSV).

File S9. Parental genotype probabilities for the diallel population in File S8 (CSV).

File S10. Trait marker phenotypes for the diallel population in File S8 (CSV).

File S11. Sequence alignment and percent identity matrix for *OFP20* (DOCX).

File S12. Marker concordance between V1 DArTag and the SNP array (CSV).

File S13. Marker concordance between V2 DArTag and the SNP array (CSV).

## AUTHOR CONTRIBUTIONS

**Jeffrey B. Endelman**: Conceptualization, Resources, Investigation, Formal analysis, Software, Supervision, Writing – original draft. **Moctar Kante**: Conceptualization, Resources, Investigation, Formal analysis, Writing – original draft. **Hannele Lindqvist-Kreuze**: Conceptualization, Resources, Supervision. **Andrzej Kilian**: Methodology. **Laura M. Shannon:** Conceptualization, Supervision. **Maria V. Caraza-Harter:** Resources. **Brieanne Vaillancourt:** Formal analysis, Data curation. **Kathrine Mailloux:** Investigation, Resources. **John P. Hamilton:** Formal analysis. **C. Robin Buell:** Conceptualization, Supervision. **All authors**: Writing – review & editing.

## ACKNOWLEDGMENTS

Development of the V1 DArTag panel was supported by the CGIAR Excellence in Breeding Platform and Crops to End Hunger Initiative. The USDA National Institute of Food & Agriculture (NIFA) Award 2019-51181-30021 supported development of the V2 DArTag panel, with additional support from PepsiCo for the development of genomic resources to validate markers. Genotyping of UW-Madison potato breeding lines was supported by USDA NIFA Awards 2020-51181-32156 and 2021-34141-35447. We thank D. DeKoeyer for suggesting the marker for *Sen3* resistance.

## DATA AVAILABILITY STATEMENT

Supplemental Files S1 – S13, which contain the marker and pedigree data needed to reproduce the results of this study, will be available from the Dryad Digital Repository at https://doi.org/10.5061/dryad.8pk0p2nw4 upon publication. Upon manuscript acceptance, PacBio HiFi sequencing data will be available via the NCBI Sequence Read Archive under BioSamples SAMN38982152, SAMN38982165, SAMN38982166, SAMN38982167, and SAMN38982169, and Illumina sequencing data will be available via the NCBI Sequence Read Archive under BioSamples SAMN39419651, SAMN39670896, SAMN39670897, and SAMN39670898.

## Supplemental Figures and Tables

Endelman *et al*. Targeted genotyping-by-sequencing of potato and data analysis with R/polyBreedR

**Figure S1.**
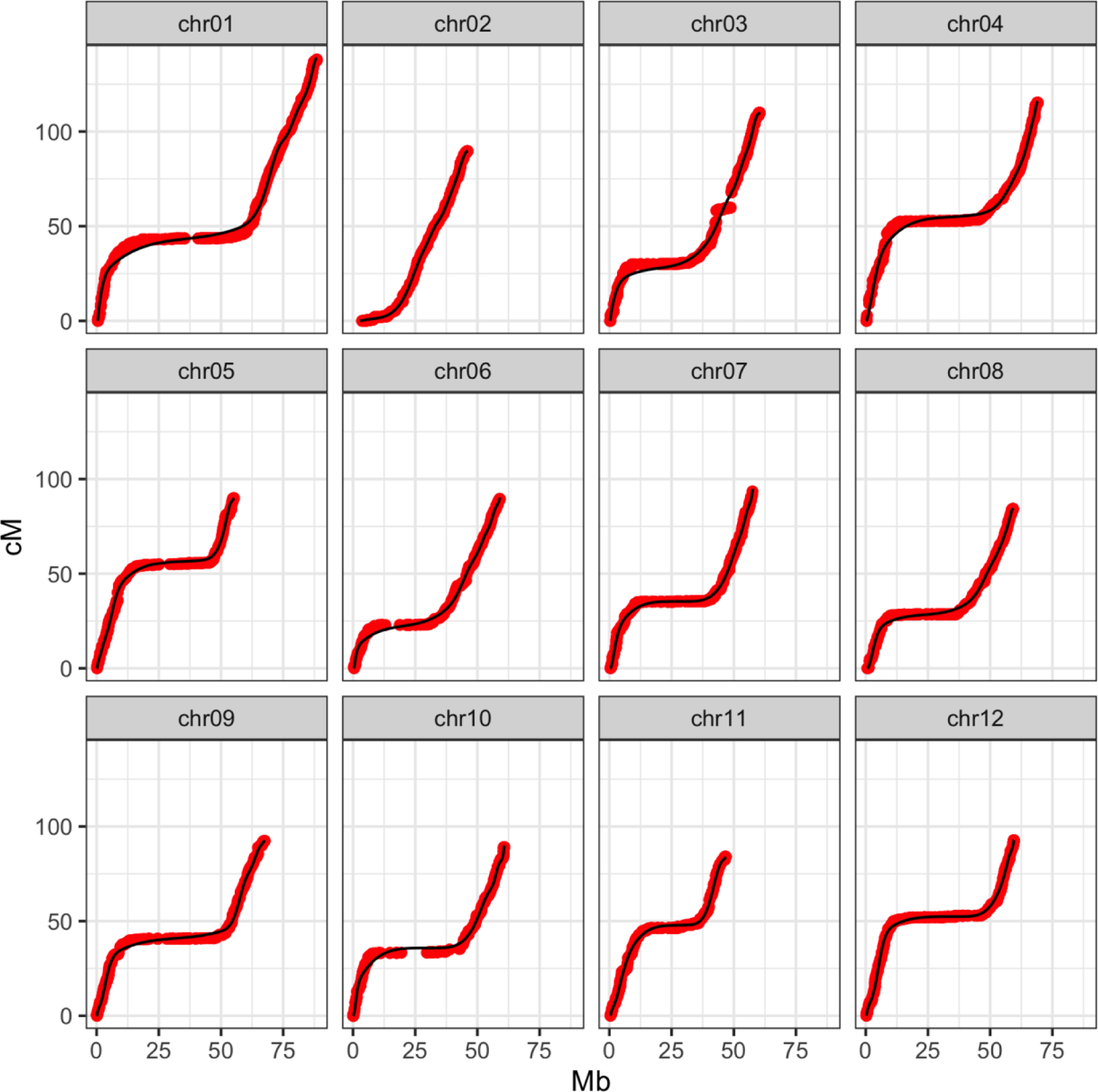
Marey Map of the potato genome. Horizontal axis is the DMv6.1 reference genome position (Pham et al., 2020).

**Figure S2.**
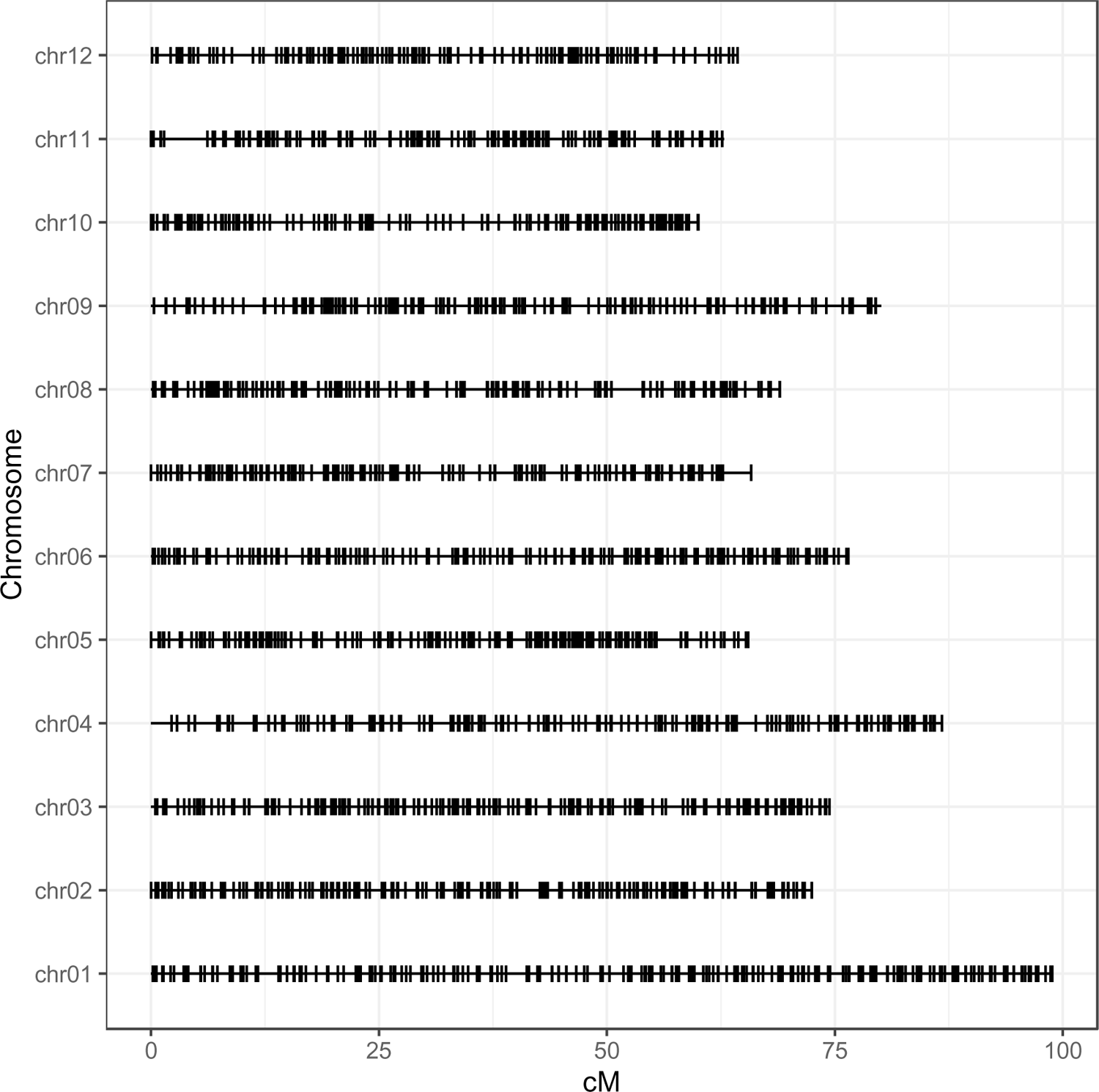
Distribution of the 2501 genomic markers for V1 DArTag.

**Figure S3.**
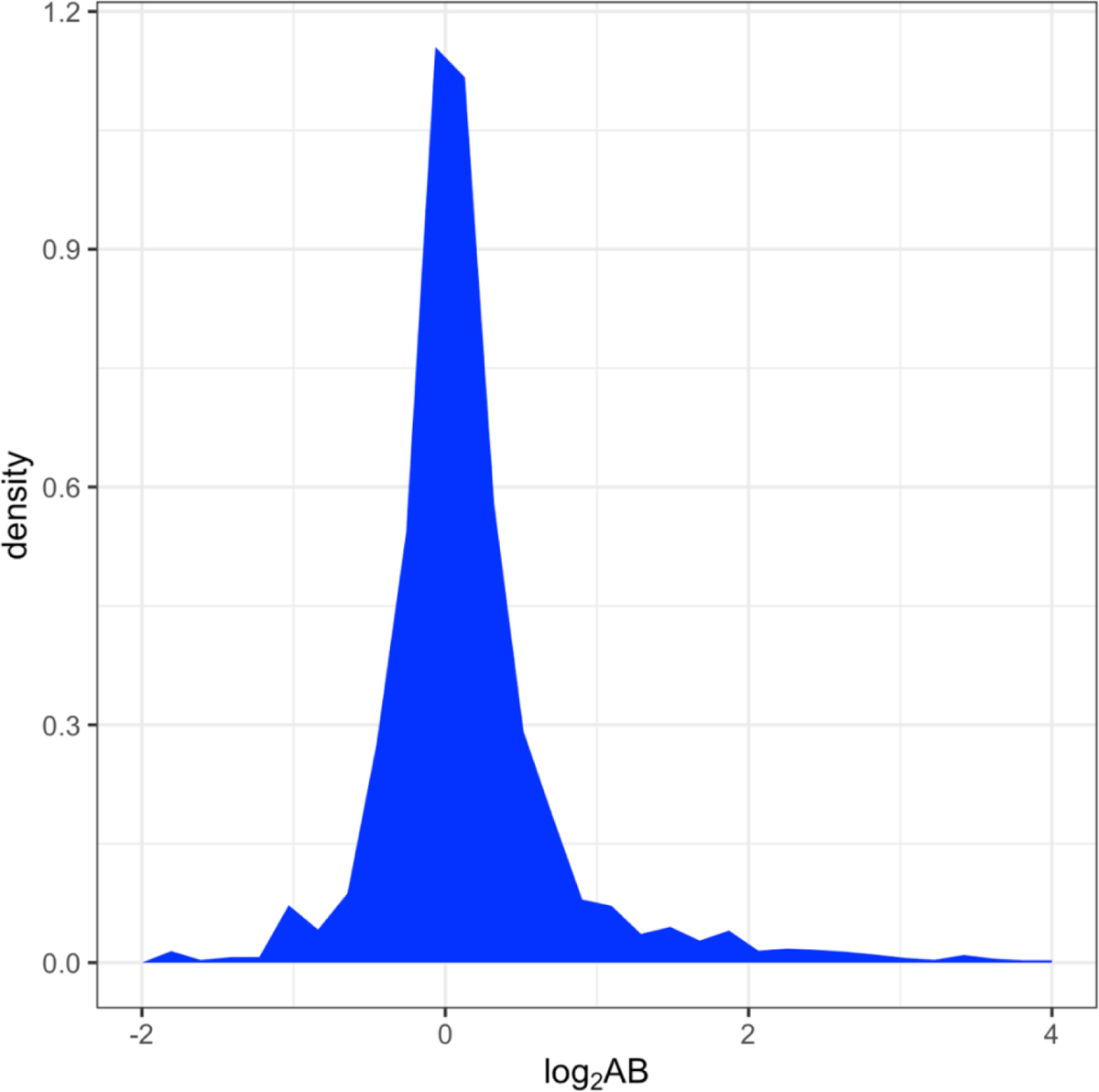
Distribution of allele bias (AB) estimates, where AB=1 indicates no bias, and values greater than 1 indicate bias toward the REF allele

**Figure S4.**
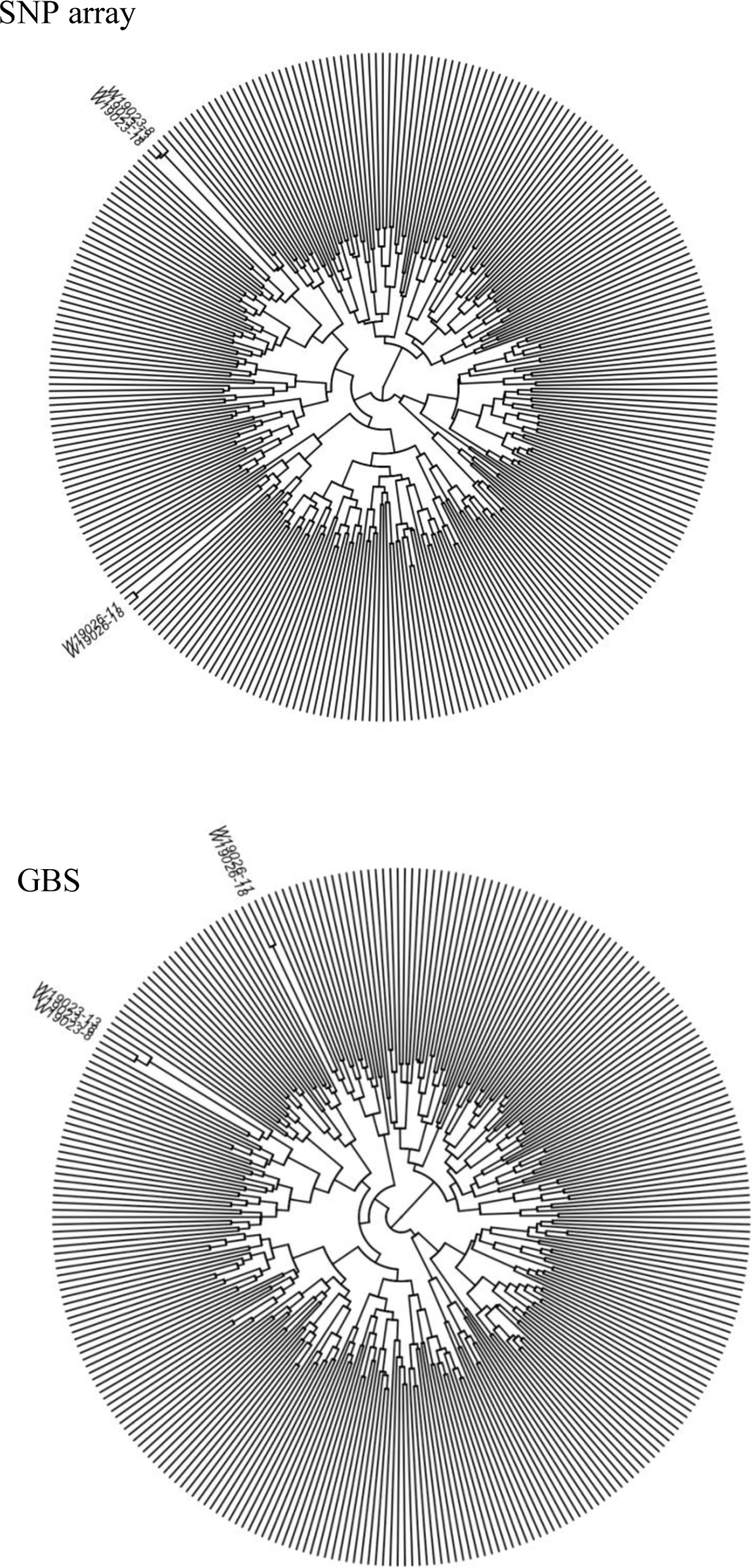
Hierarchical clustering based on SNP array (top) or GBS (bottom) data. Both platforms identified two groups of genetically identical clones, one pair and one threesome, originating from the same F1 populations

**Figure S5.**
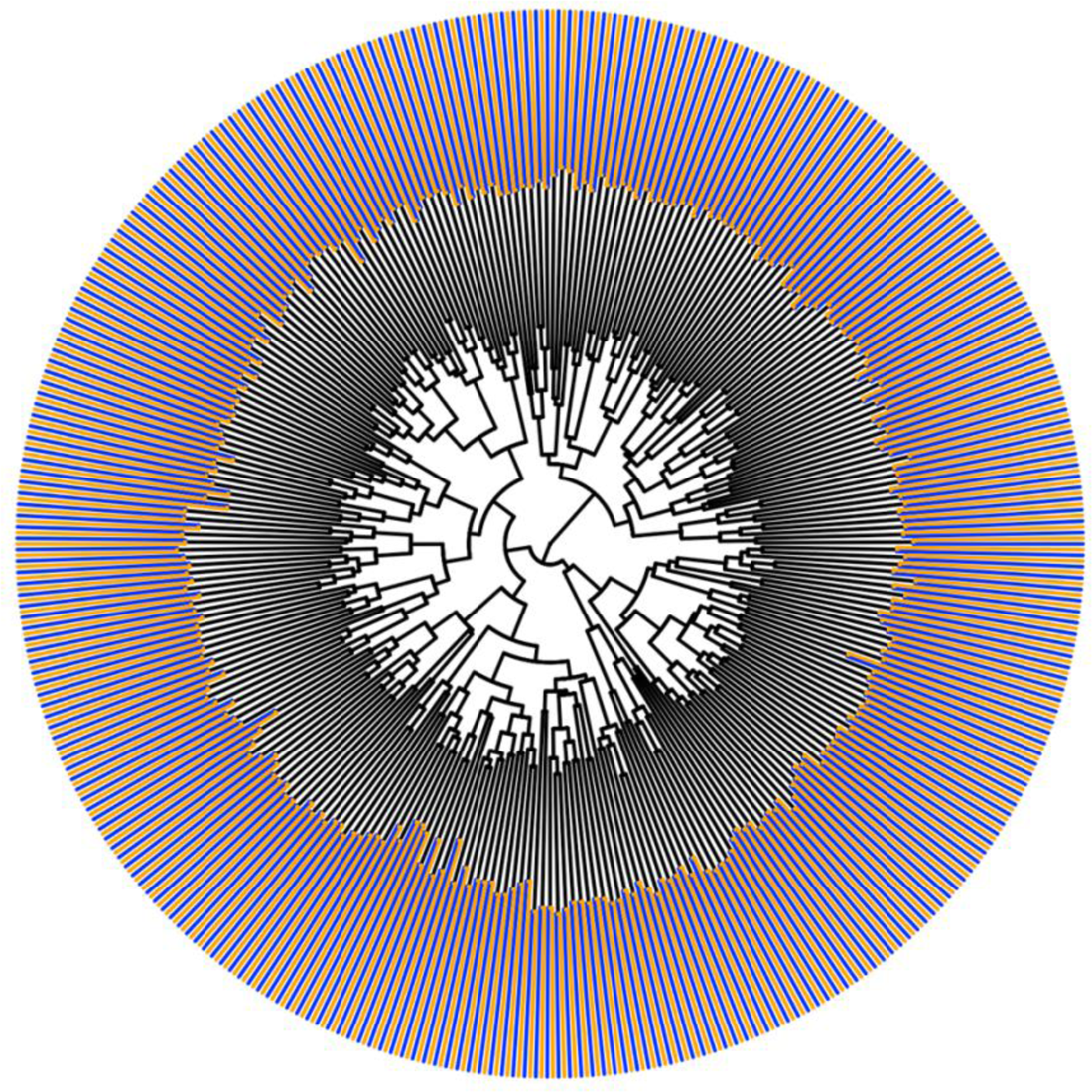
Joint clustering of SNP array (blue) and GBS (orange) samples. The two marker profiles for every clone were paired.

**Figure S6.**
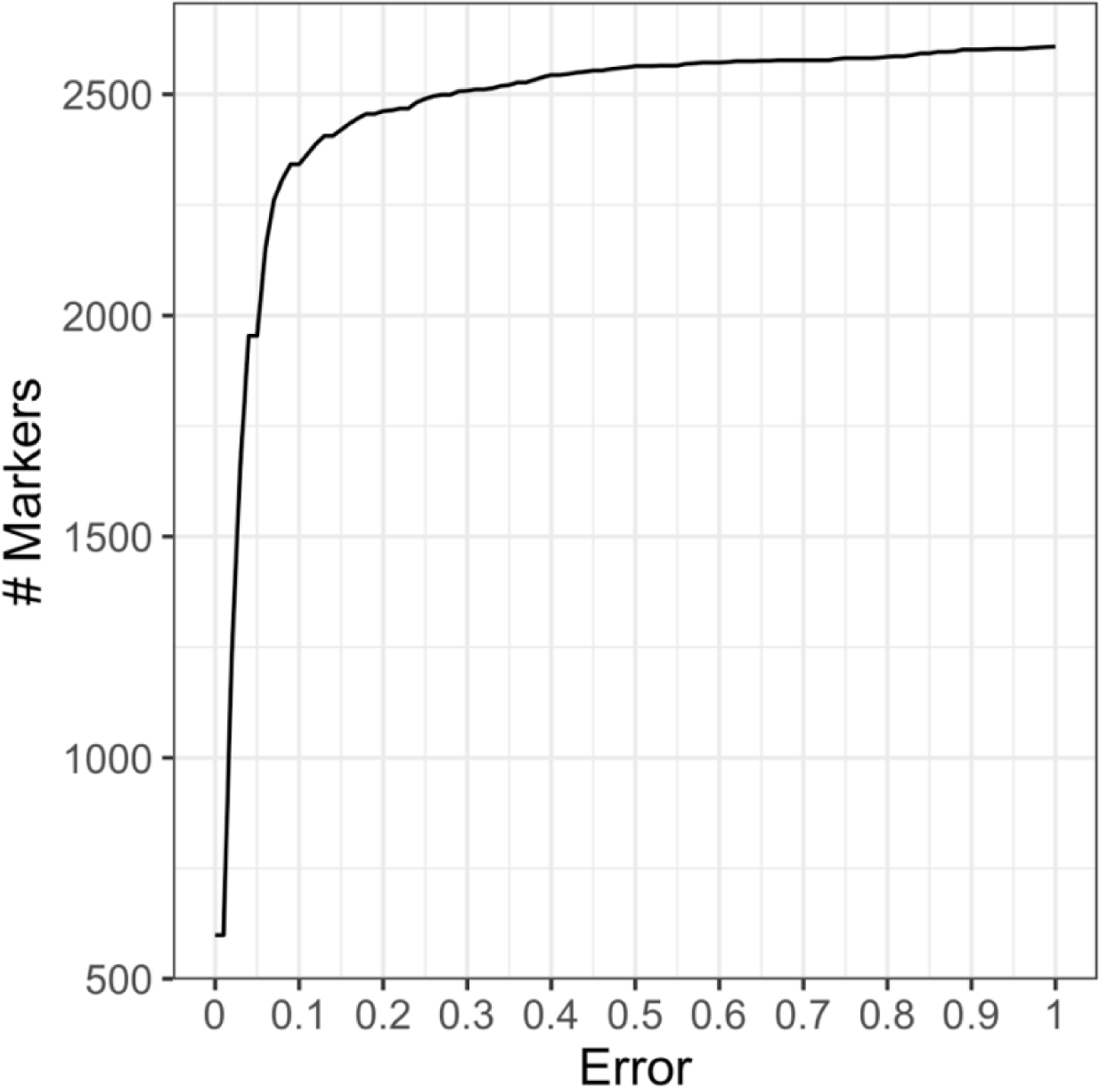
Empirical cumulative distribution for 2x classification error for 2608 common markers between the V4 SNP array and V2 DArTag.

**Figure S7.**
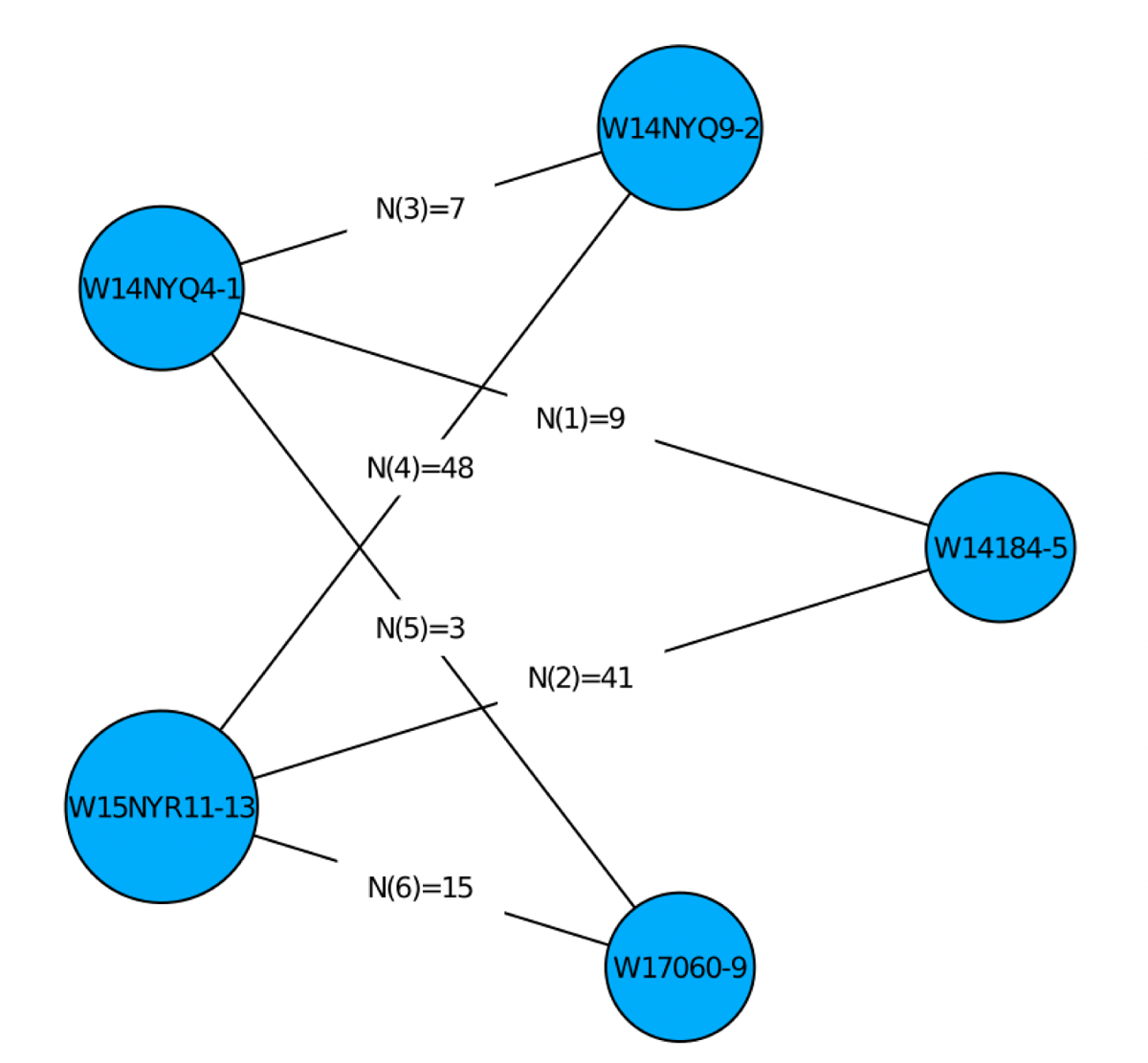
Five-parent partial diallel. Graphical output from PolyOrigin shows the number of progeny per biparental F1 population.

**Figure S8.**
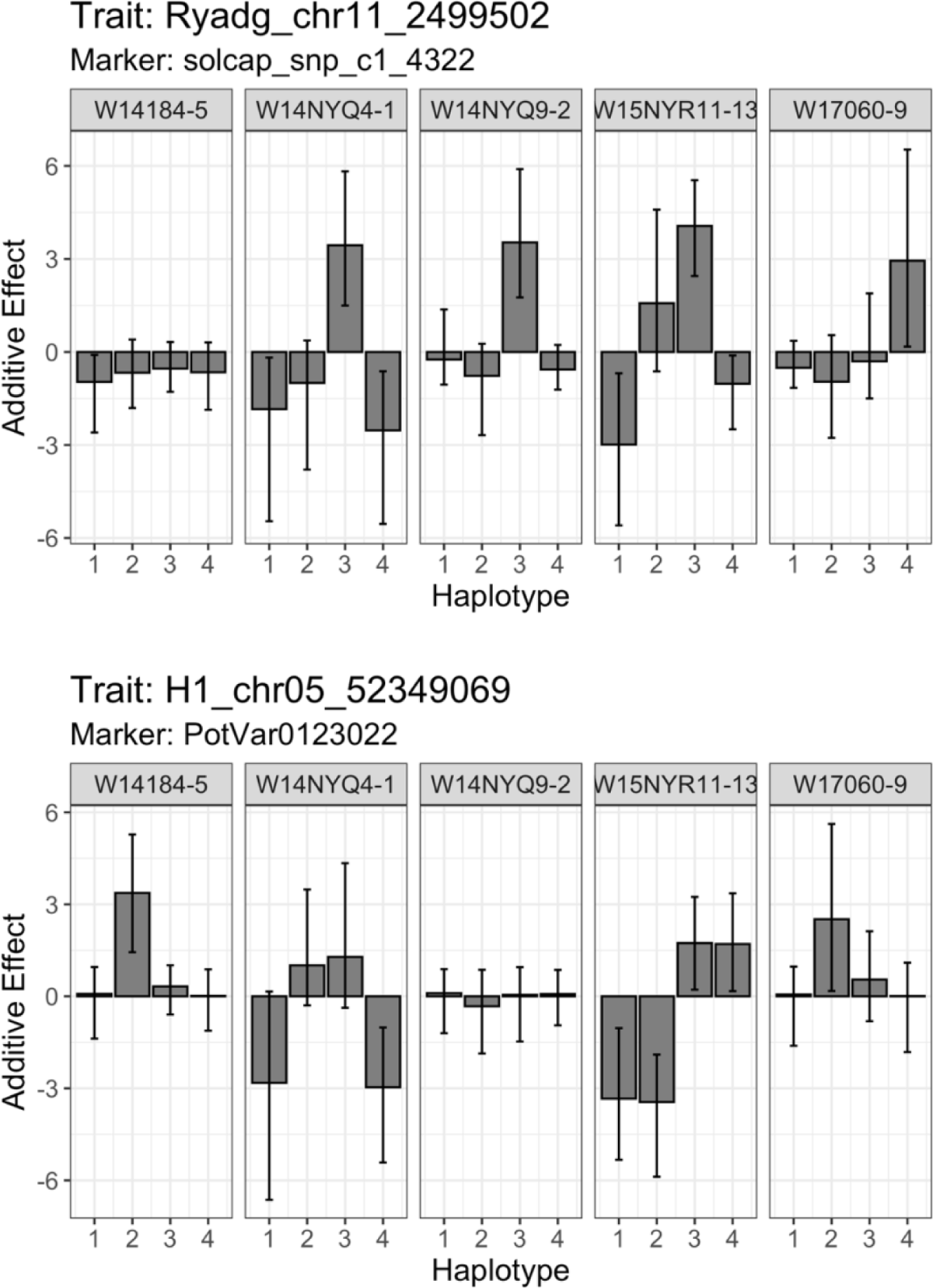
Additive effect estimates for parental haplotypes in the five-parent partial diallel. Positive values indicate presence of the R gene. From left to right, the result indicates the parental dosage of Ryadg is 0, 1, 1. 2, 1, and for H1 the parental dosage is 1, 2, 0, 2, 1. Parents W14NYQ9-2 and W15NYR11-13 were included in the V2 DArTag submission, and their genotype calls agree with these predictions (FileS3).

**Table S1.** Comparison of KASP (rows) vs. V1 DArTag (columns) markers targeting *Ry_adg_* (snpST00073).

|  | **Absent** | **Present** |
| --- | --- | --- |
| **Absent** | 7 | 1 |
| **Present** | 1 | 84 |

**Table S2.**
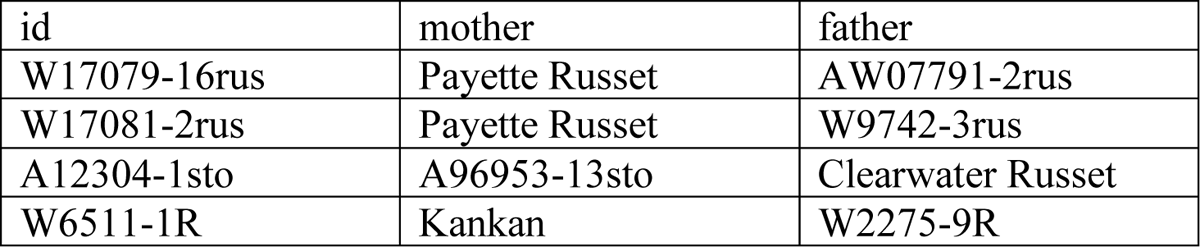
Positive samples for marker Rysto_chr12_2352742 in V2 DArTag.

| id | mother | father |
| --- | --- | --- |
| W17079-16rus | Payette Russet | AW07791-2rus |
| W17081-2rus | Payette Russet | W9742-3rus |
| A12304-1sto | A96953-13sto | Clearwater Russet |
| W6511-1R | Kankan | W2275-9R |

**Table S3.** Results for marker Sli_chr12_2372490 in V2 DArTag.

| id | ALT dosage |
| --- | --- |
| W9968-5-DH027 | 0 |
| CO99076-6R-DH002 | 0 |
| CO99076-6R-DH033 | 0 |
| W14NYQ29-5-DH024 | 0 |
| W14NYQ9-2-DH119 | 0 |
| W14NYQ9-2-DH132 | 0 |
| W8890-1R-DH003 | 0 |
| W9968-5-DH084 | 0 |
| W9968-5-DH022 | 0 |
| W13069-5-DH088 | 0 |
| RioColorado-DH003 | 0 |
| W9968-5-DH151 | 0 |
| W12078-76-DH352 | 0 |
| W12078-76-DH099 | 0 |
| W13058-4-DH002 | 0 |
| W14NYQ9-2-DH137 | 0 |
| W14NYQ9-2-DH146 | 0 |
| RioColorado-DH005 | 1 |
| W15263-50R-DH001 | 1 |
| W15263-50R-DH011 | 1 |
| W8890-1R-DH002 | 1 |
| W9426-3R-DH005 | 1 |
| W9426-3R-DH037 | 1 |
| W2x150-24 | 1 |
| W2x113-3 | 1 |
| W2x001-22-45 | 2 |
| W2x082-(14/20)-13 | 2 |
| W2x082-(14/20)-13-2 | 2 |

## Supplemental Methods

Endelman et al. (2024)

This document shows how the main figures and tables were generated on a MacOS system. R packages polyBreedR and diaQTL can be installed from their GitHub sites. Other R packages are available on CRAN. The impute_LA function utilizes PolyOrigin, which is available from its GitHub site and requires installation of Julia. Command line calls to bcftools are used to manipulate VCF files.

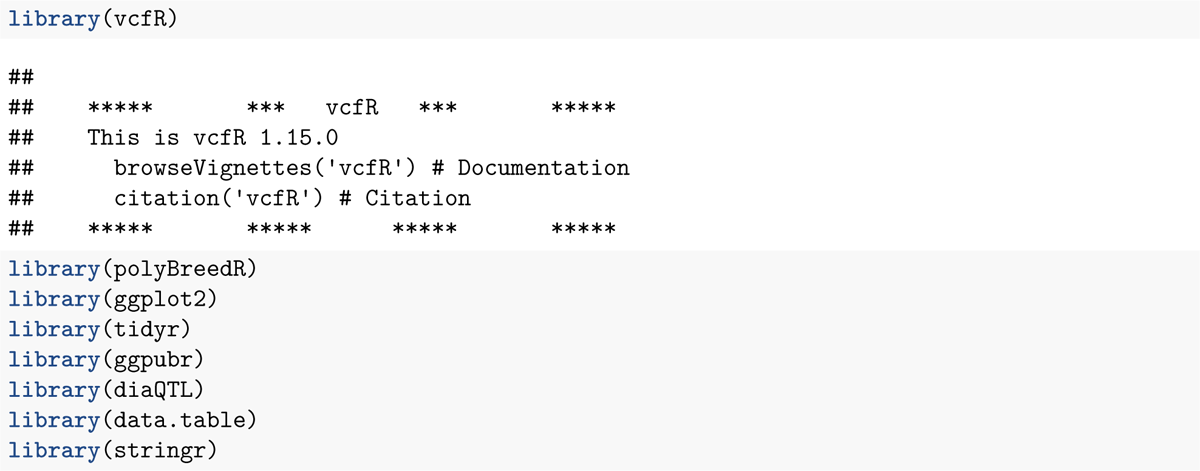

**Figure 1.**
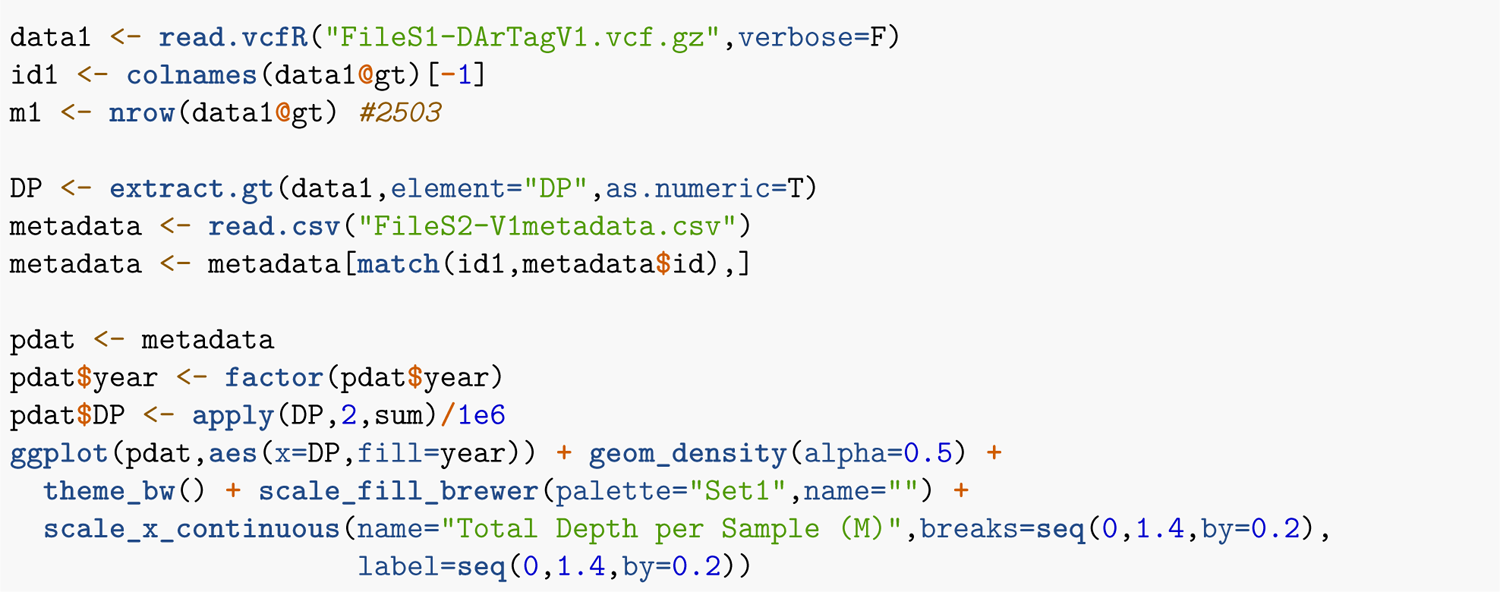
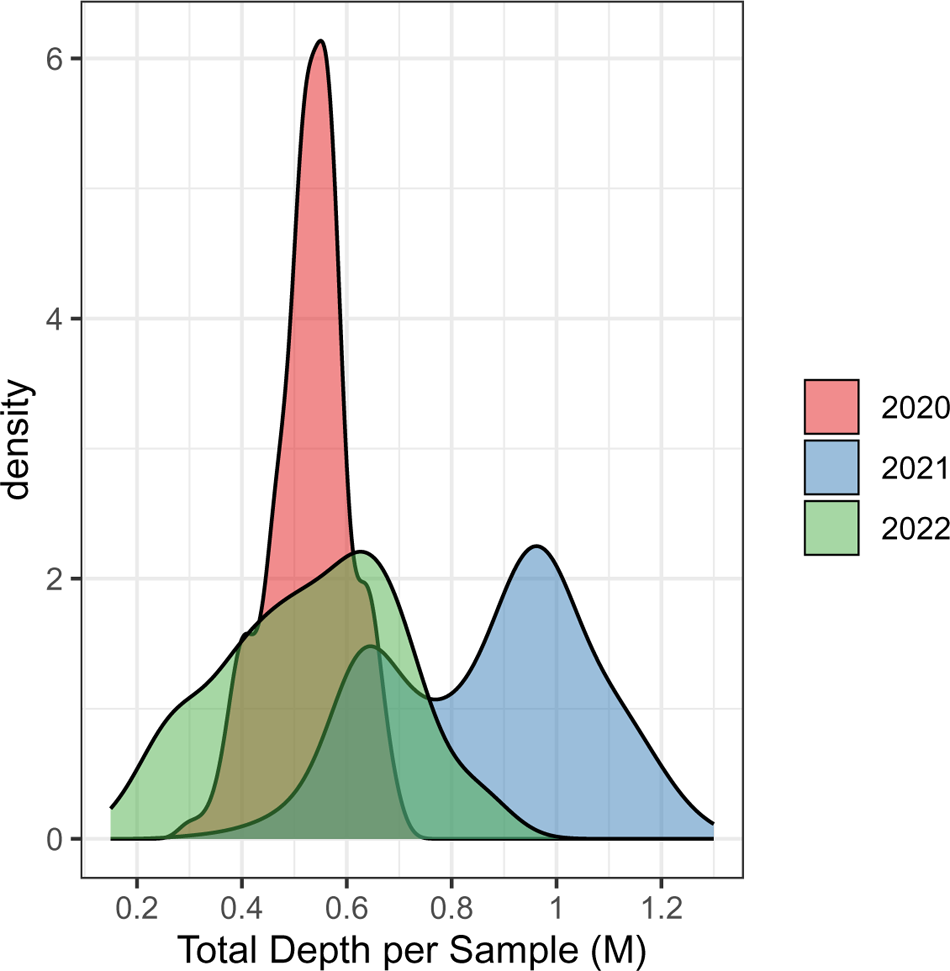

**Figure 2.**
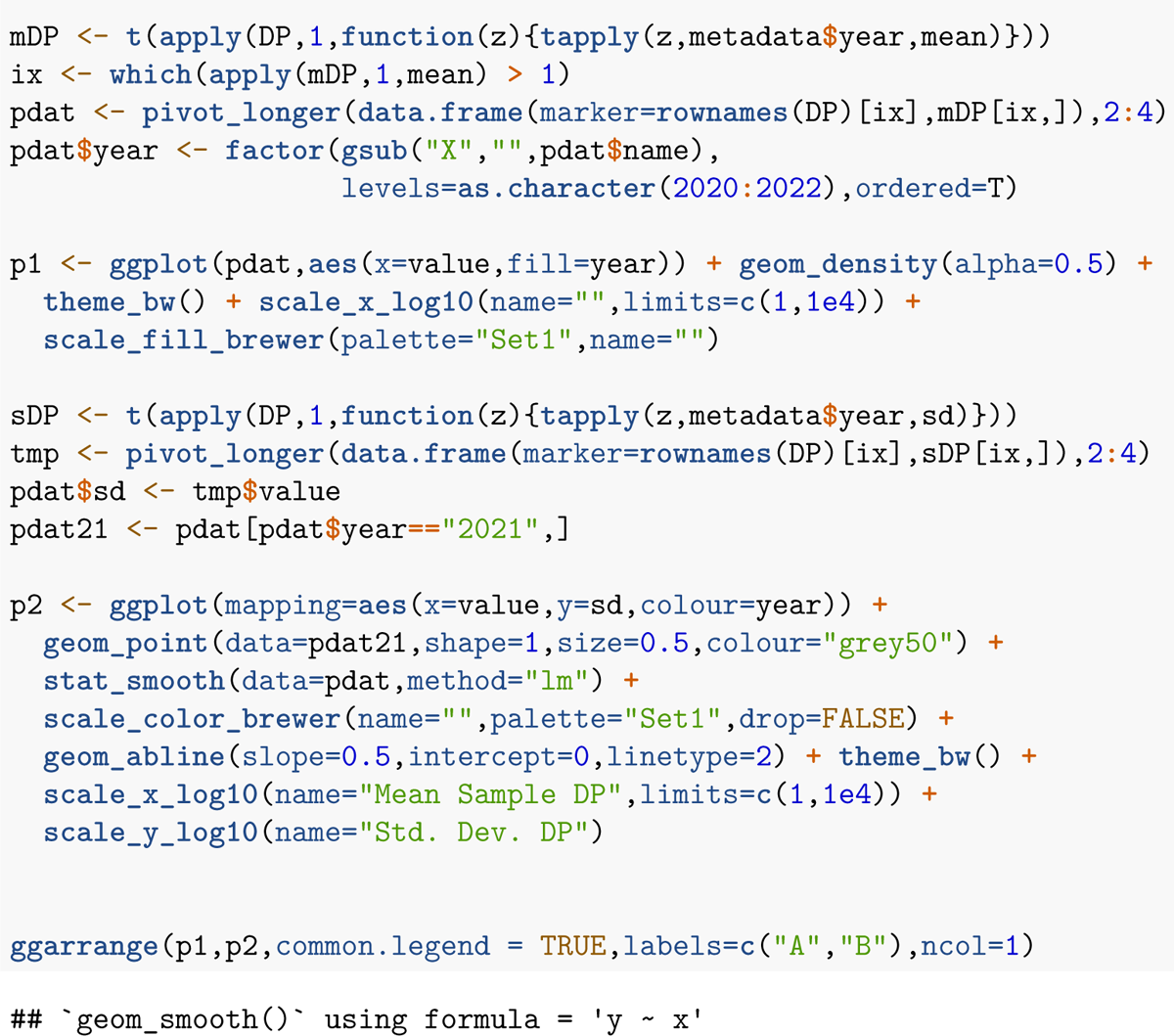
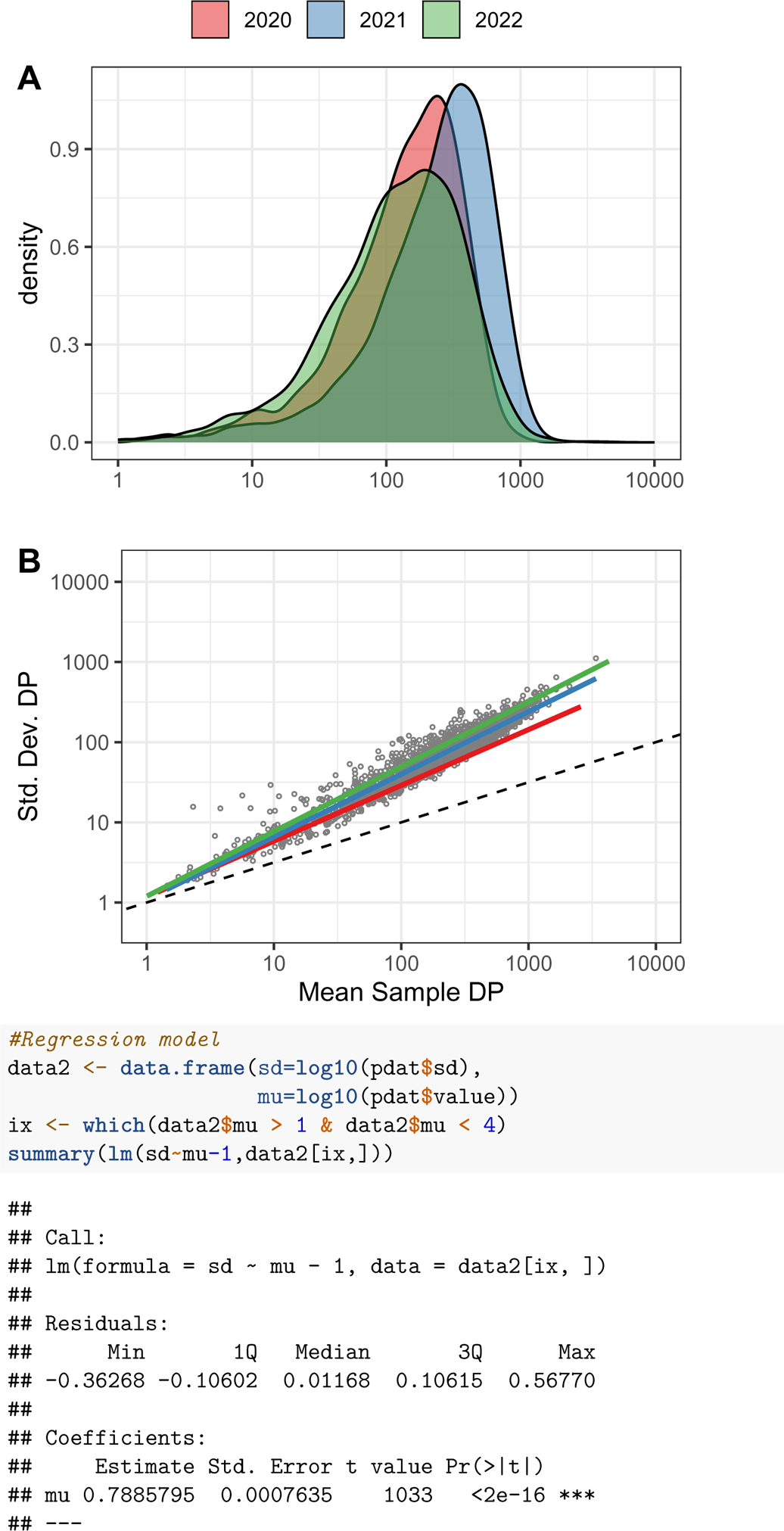
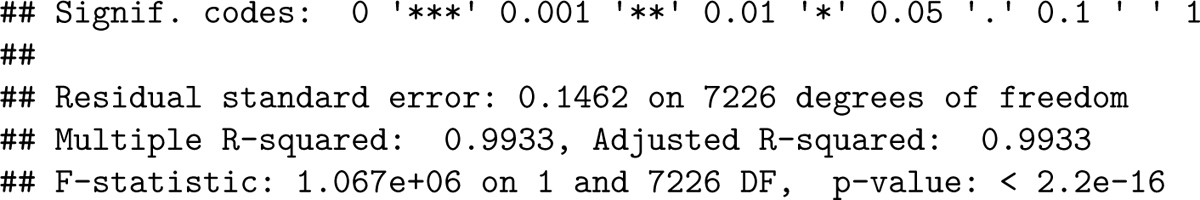

**Figure 3.**
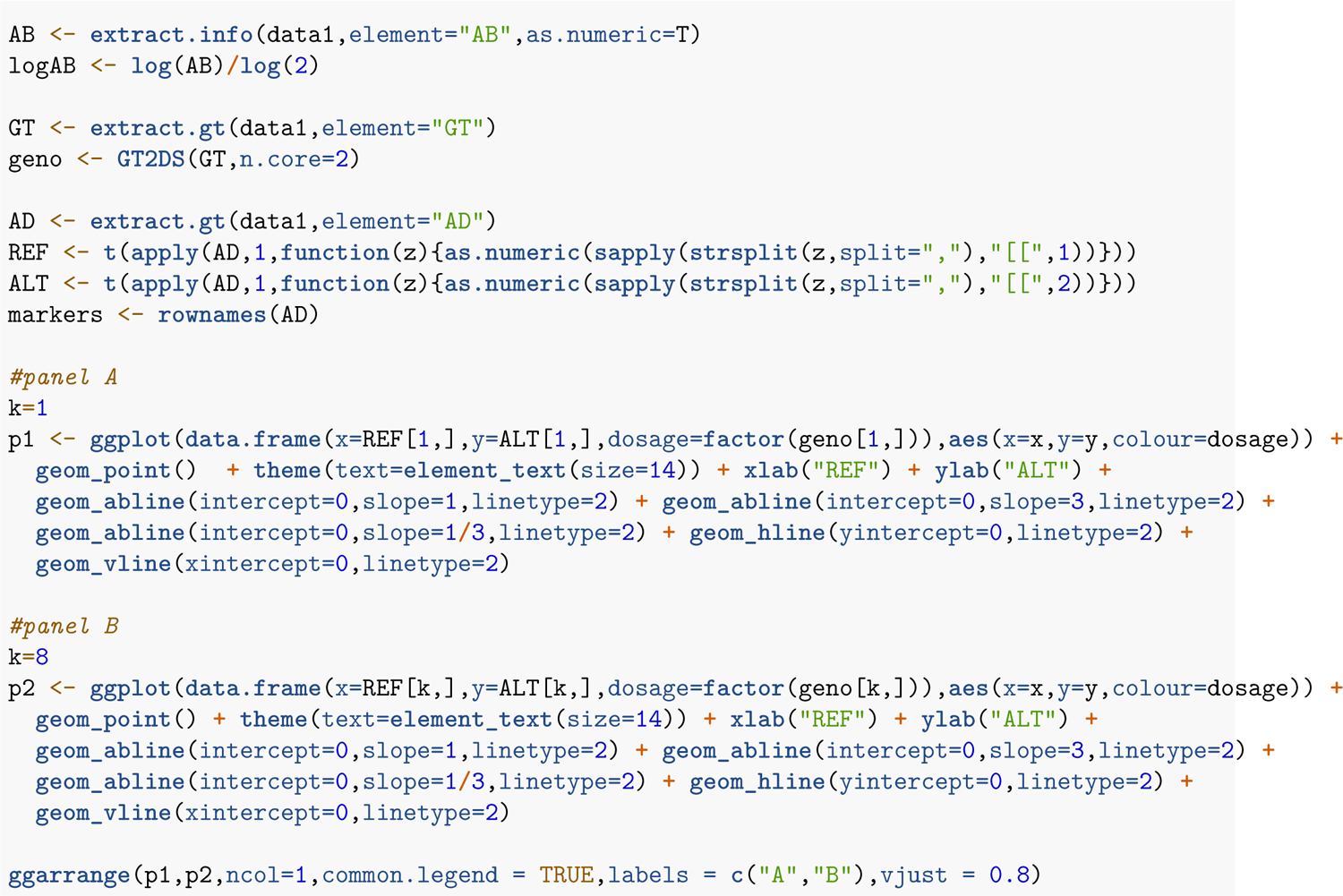
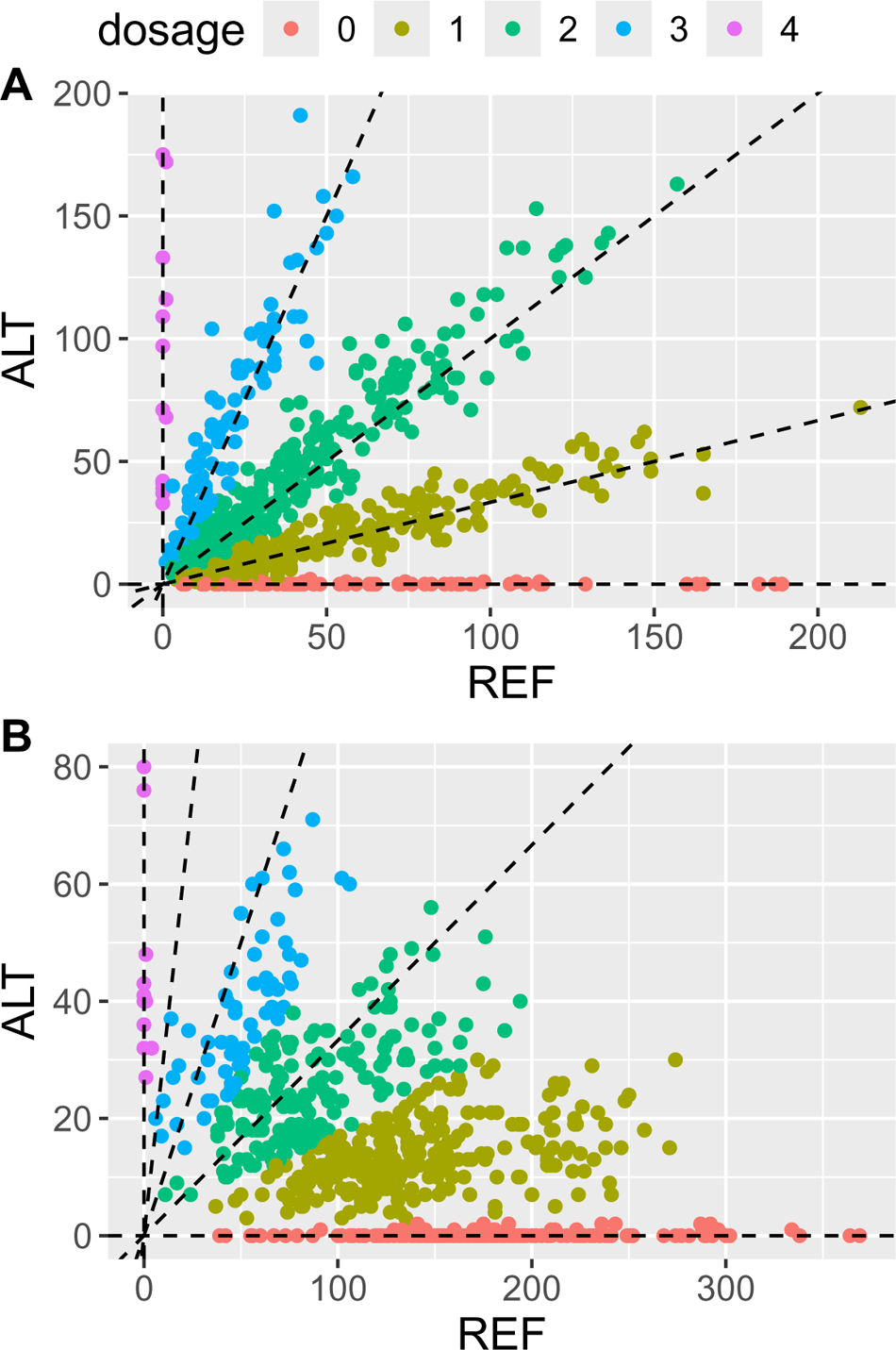

**Figure 4.**
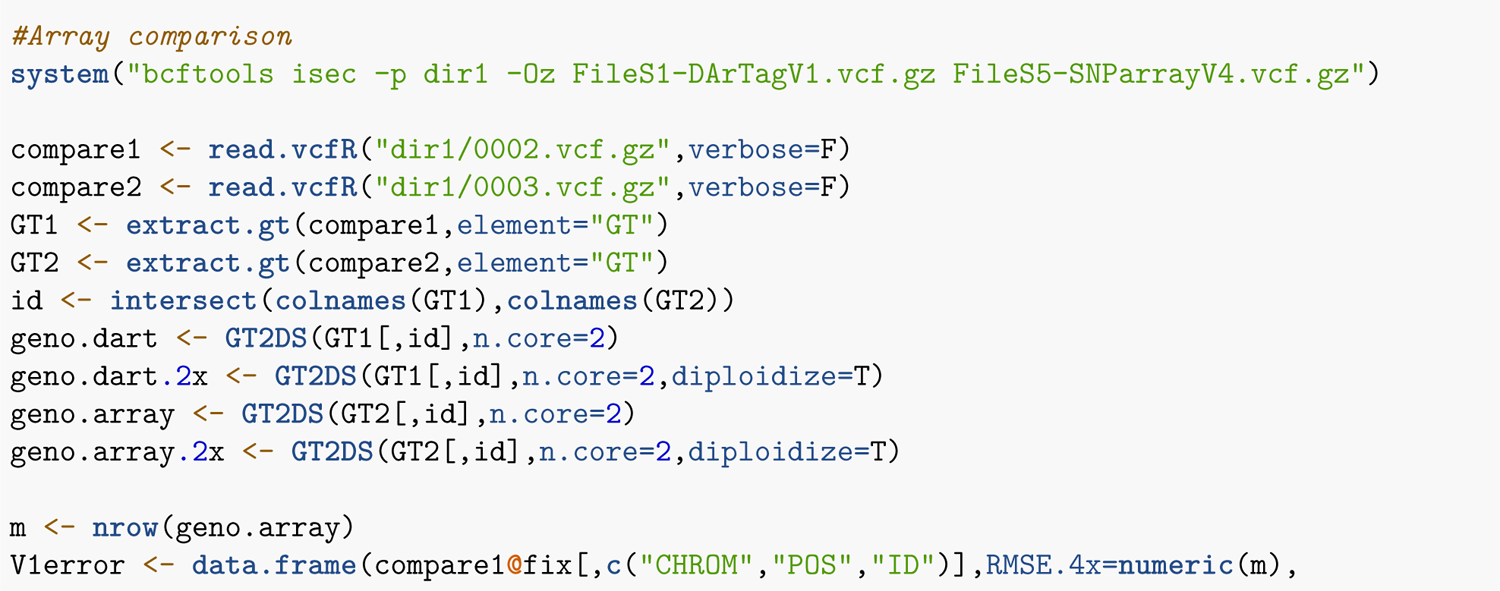
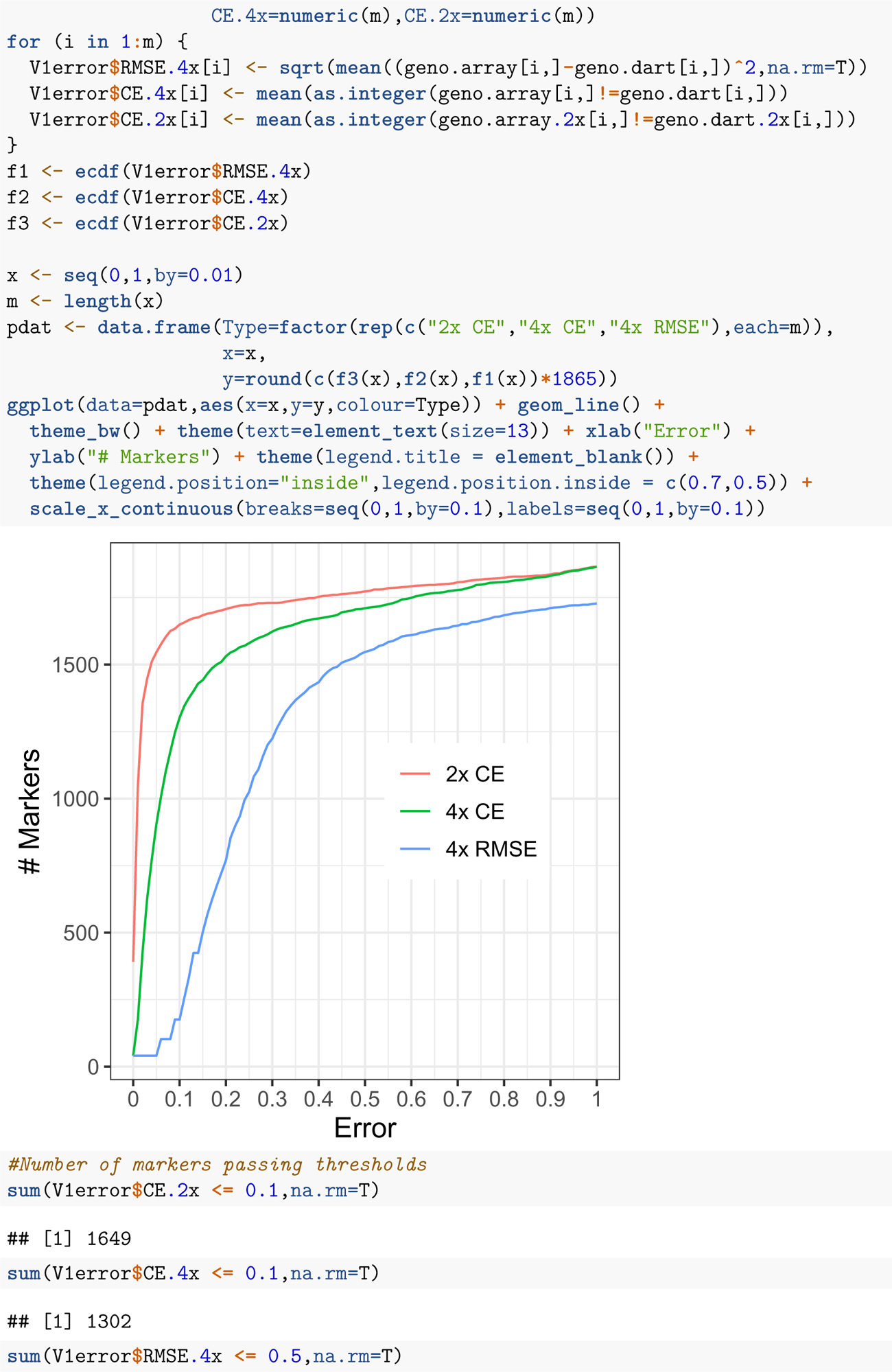
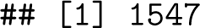

**Figure 5.**
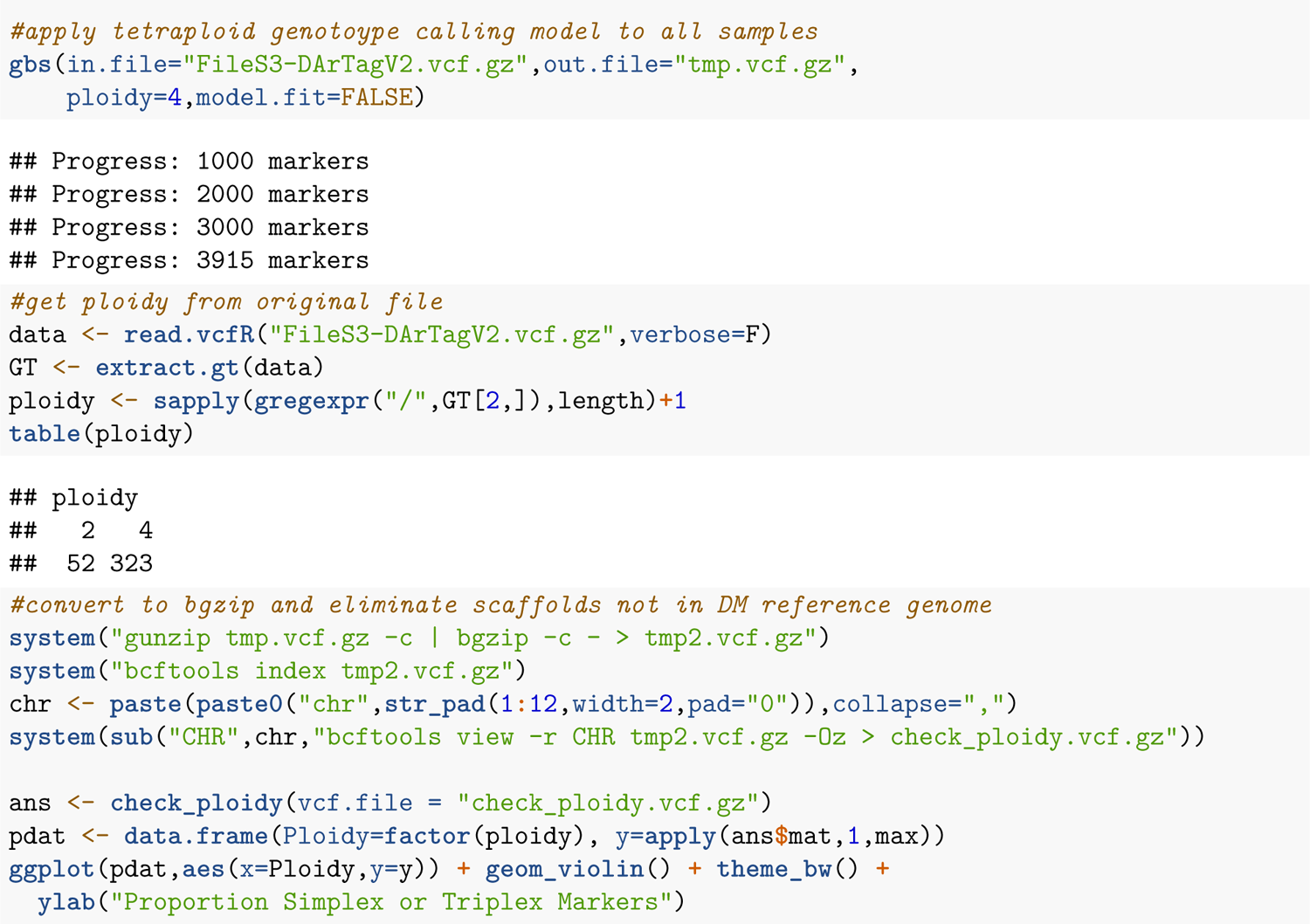
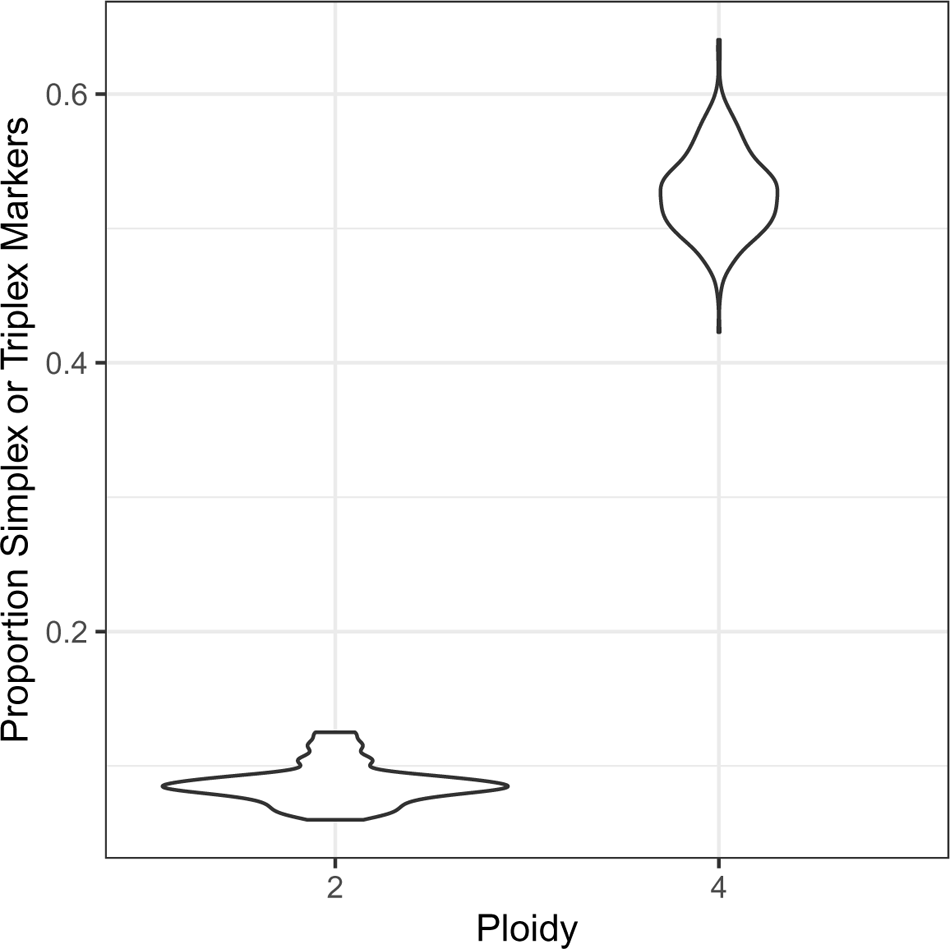

**Table 1.**
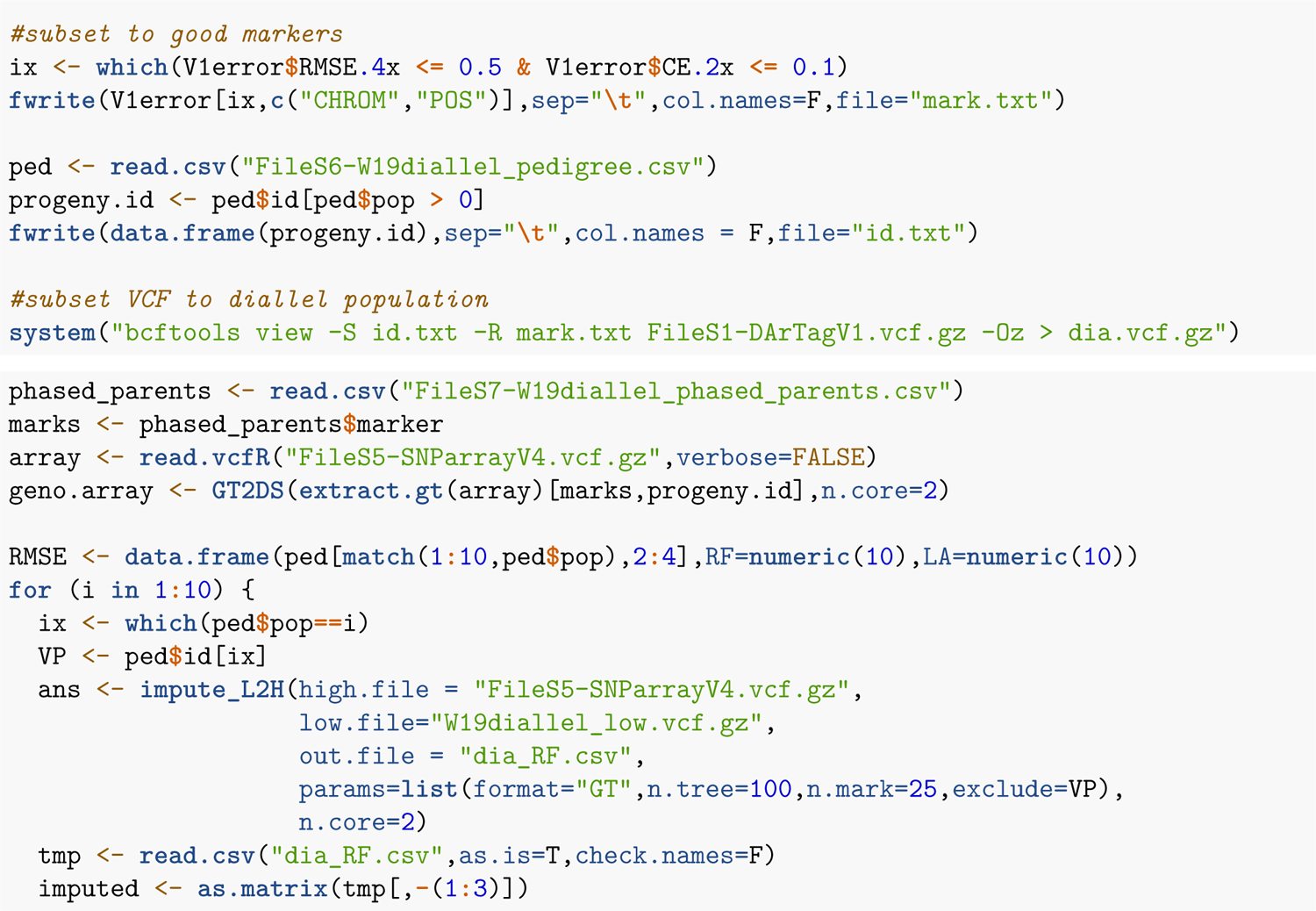

**Figure 6.**
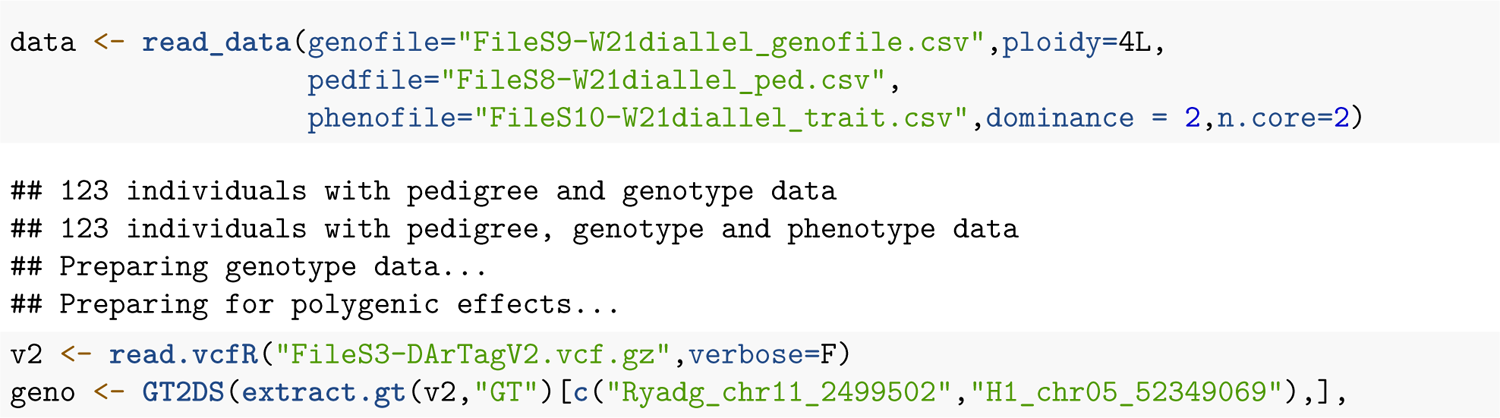
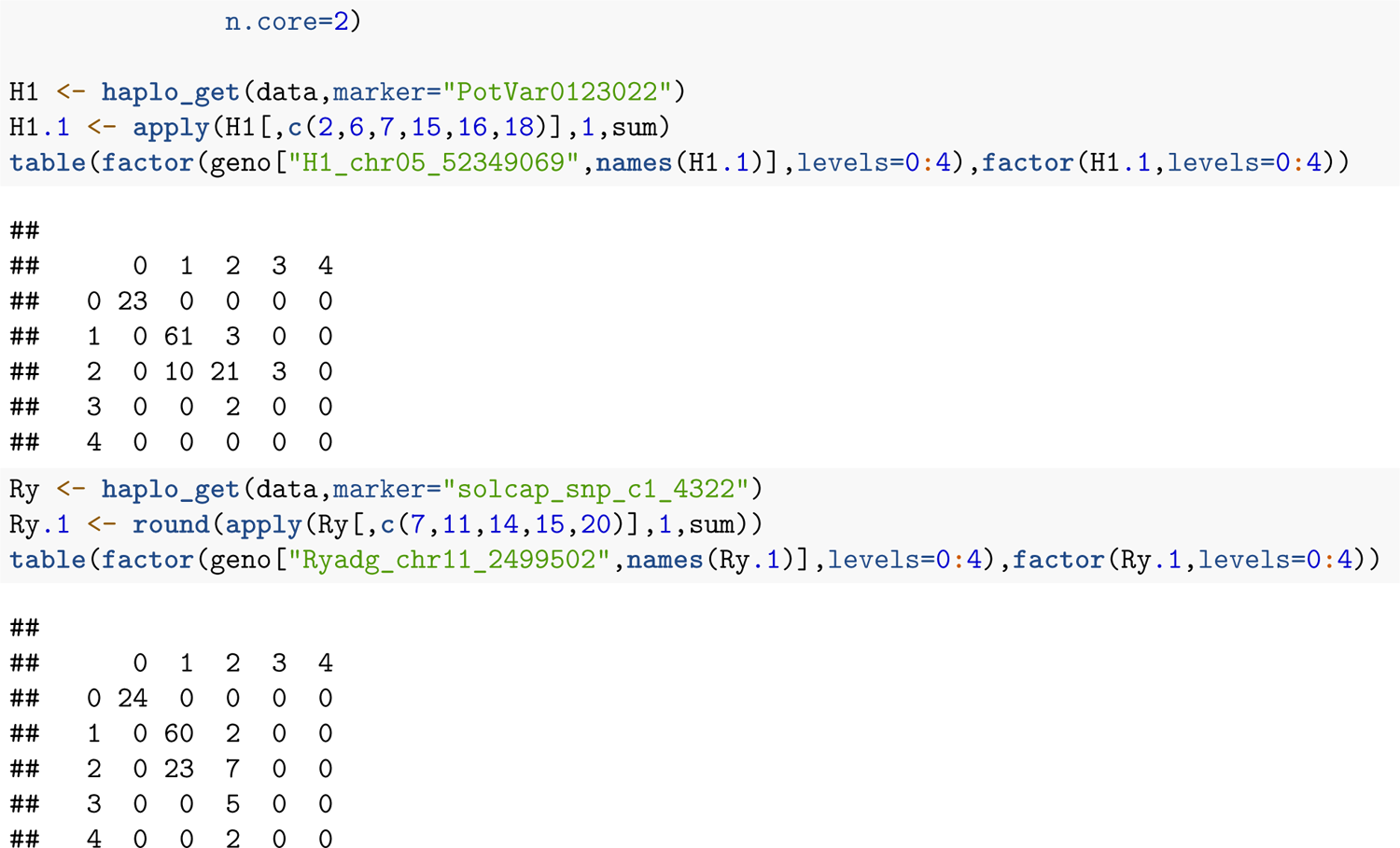

**Figure 7.**
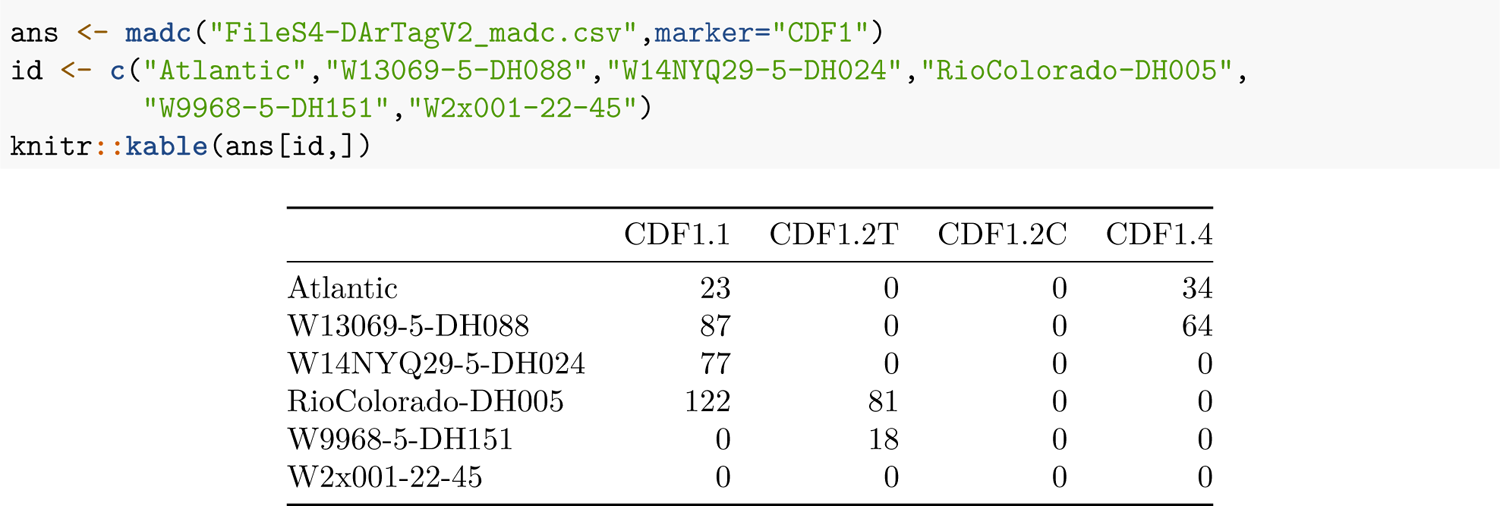

**Figure 8.**
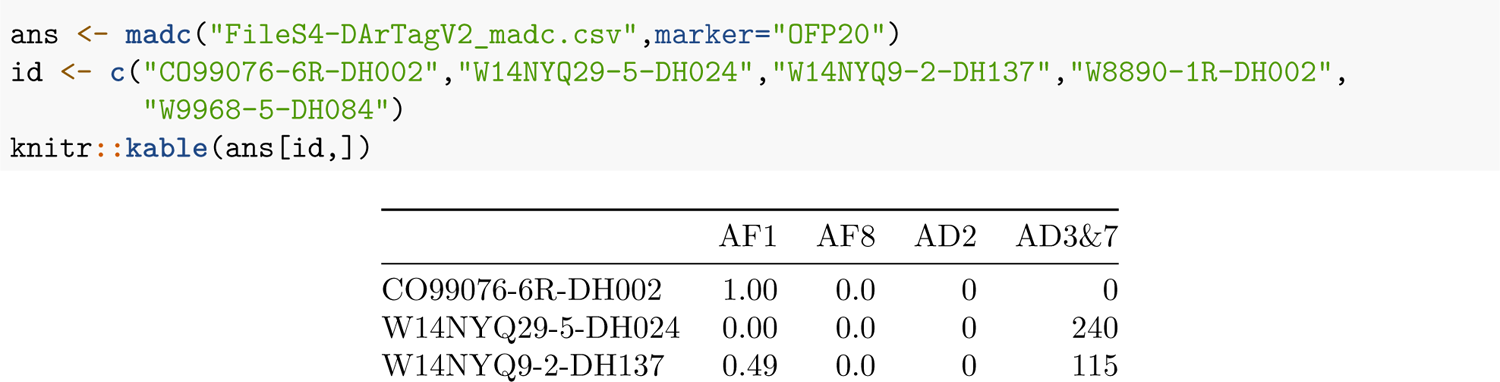
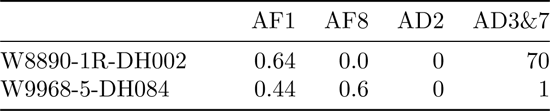

